# Effects of Hypomethylating Agents on Gene Modulation in the Leukemic Microenvironment and Disease Trajectory in a Mouse Model of AML

**DOI:** 10.1101/2024.12.01.626276

**Authors:** Nancy D. Ebelt, Suvithanandhini Loganathan, Lara C. Avsharian, Edwin R. Manuel

**Author notes:** (N.D.E.) (L.C.A); (S.L.); (E.R.M).

## Abstract

Hypomethylating agents (HMAs), such as decitabine and 5-azacytidine (AZA), are valuable treatment options for patients with acute myeloid leukemia that are ineligible for intensive chemotherapy. Despite providing significant extensions in survival when used alone or in combination, eventual relapse and resistance to HMAs are observed. The mechanisms leading to these outcomes are still not well defined and may, in part, be due to a focus on leukemic populations with limited information on the effects of HMAs on non-leukemic cells in the blood and other tissue compartments. In this study, we elucidated effects on immune-related gene expression in non-leukemic blood cells and the spleen during AZA treatment in leukemia-challenged mice. We observed significant changes in pathways regulating adhesion, thrombosis, and angiogenesis as well as a dichotomy in extramedullary disease sites that manifests during relapse. We also identify several genes that may contribute to the anti-leukemic activity of AZA in blood and spleen. Overall, this work has identified novel gene targets and pathways that could be further modulated to augment efficacy of HMA treatment.

## INTRODUCTION

Patients with acute myeloid leukemia (AML) and other myelodysplastic syndromes (MDS) are frequently ineligible for standard of care chemotherapy and hematopoietic stem cell transplantation (HCST) due to extensive co-morbidities^1,2^. In these cases, hypomethylating agents (HMA) such as azacytidine or decitabine are well tolerated and show overall response rates as high as 50%^3,4^. The correlation of BCL-2 expression in AML patients with poorer overall survival^5,6^ and the pre-clinical and clinical discoveries of synergy between BCL-2 inhibition and HMAs^7,8^ have resulted in the approval of 5-azacytidine (AZA) and venetoclax (small molecule inhibitor or BCL-2) by the FDA in 2017. This combination therapy increases the overall response rate to 66% and overall survival significantly (median follow up 20.5 months)^9^. Despite these advances, many patients that respond to the combination treatment will relapse within 3+ years, with overall survival reaching only 35-40%^10^, similar to azacytidine monotherapy (2-year survival ~50%)^11^. Determining the direct, cause-effect relationship between HMAs and anti-leukemia efficacy is necessary to determine additional, rational treatment combinations to increase long-term remission for AML patients.

Studies of the precise mechanisms of HMA action have involved methylation array, gene expression array but only of sorted blasts or blast-heavy bone marrow aspirates, despite the fact that HMA efficacy has a large immune component involving the action of non-cancerous white blood cells^12–20^. These studies are highly conflicting with regard to methylation and gene expression signatures found, and although some signatures had prognostic significance, few were able to help narrow down a precise mechanism for HMA efficacy^21–25^, demonstrating the need for studies of other AML microenvironments and constituents in addition to the blasts themselves. After AML remission from chemotherapy, increased levels of neutrophils and platelets as determined by complete blood counts were shown to independently predict longer relapse free survival in a large patient cohort^26^. The time to recovery of platelets and neutrophils after induction chemotherapy also has prognostic value with faster recovery predicting longer relapse-free survival ^27^. In blood plasma, high levels of fibrinogen were found to be an independent indicator for worse overall survival^28,29^. It is inconclusive whether these parameters have any prognostic value after treatment with HMAs.

For this manuscript we have focused on the mechanisms of AZA efficacy in the immune microenvironments of peripheral blood and spleen that may harbor insights into the causes of eventual relapse and altered disease outcomes.

## RESULTS

### Hematologic Profiles in Blood of AZA-treated, Leukemic Mice Resemble Those in Healthy Mice

In our previous study, a single round of AZA treatments (three treatments per week) in the C1498 model only resulted in a modest increase in survival, indicating that mice never reached complete remission and a majority of blasts were still present ^12^. To ensure full penetrance of AZA treatment to more resistant clones, we continuously treated mice thrice weekly and survival was recorded. C1498-implanted, AZA-treated mice (C1498-AZA) survived significantly longer than C1498-implanted, DMSO-treated mice (C1498-vehicle) (**Figure 1A**, p<0.0001, Mantel-Cox test), with C1498-vehicle mice having a median survival of 21 days and AZA mice having a median survival of 40 days. These data are consistent with human AML patients experiencing increased benefit from prolonged AZA treatment beyond initial response^30^.

**Figure 1.**
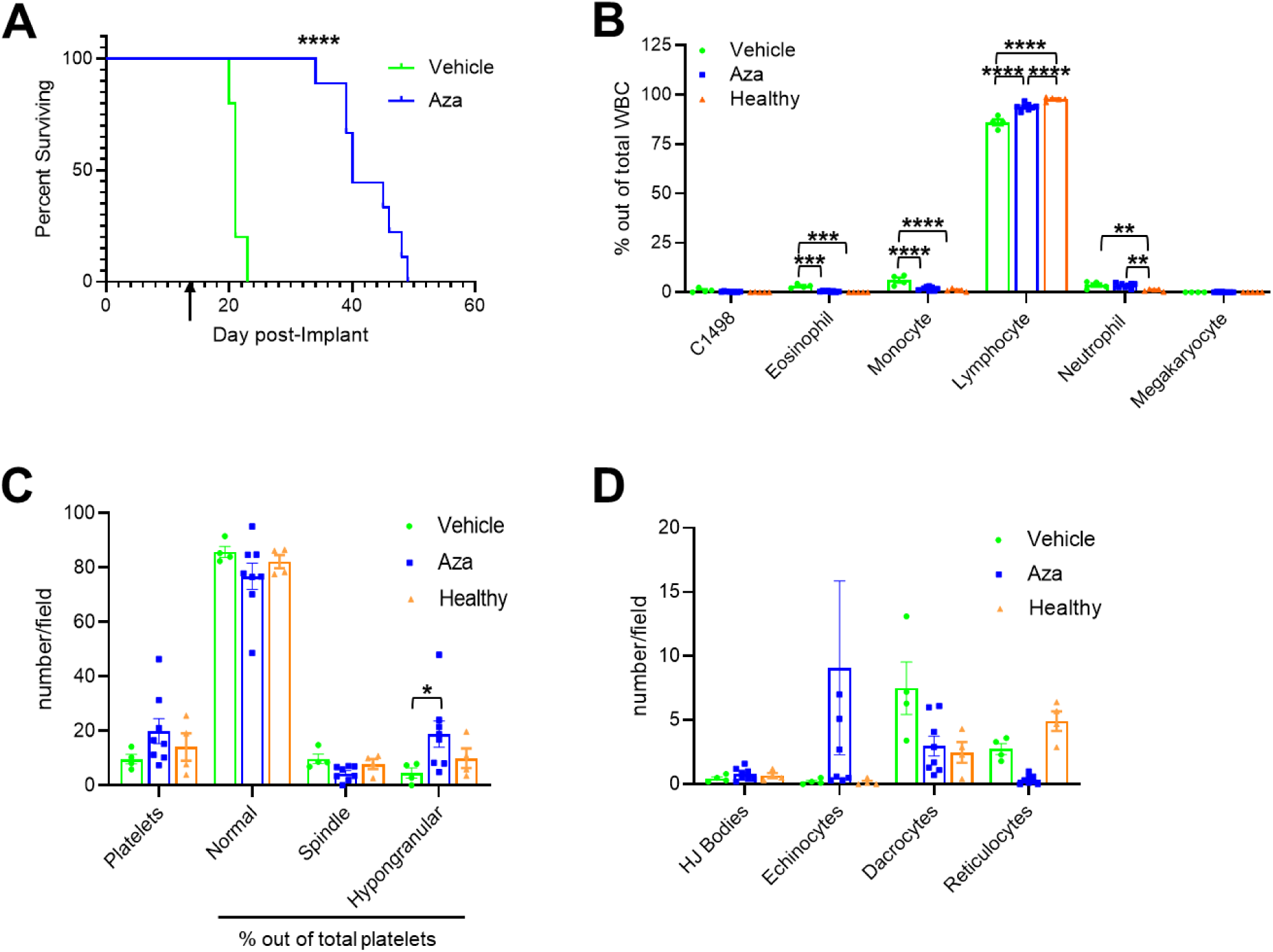
Survival and Hematologic Profiles of Leukemic Vehicle- and AZA-treated Immunocompetent Mice. C1498 leukemia cells (80,000) were implanted in C57Bl/6 mice via tail vein injection. Treatment with DMSO in diluent (Vehicle) (n=5) or 5mg/kg/mouse AZA (n=11) by intraperitoneal injection began on day 3 after implantation and continued thrice weekly until survival endpoints. **(A)** Mice were euthanized once they showed signs of terminal disease state and survival was recorded for each mouse and displayed as a Kaplan-Meier curve. Statistical significance was analyzed using the Log-rank (Mantel-Cox) test: ****p<0.0001. The arrow (↑) indicates the timepoint at which retroorbital blood was drawn to assess hematologic profiles. **(B)** Blood drawn at week 2 after treatments was smeared and stained for manual complete blood counts. Blood from non-leukemic, untreated (Healthy) mice was also included. White blood cells (WBC) were counted manually, including C1498, and are presented as a percentage of total WBC counted. **(C)** Numbers of platelets and percentages of normal, spindle-shaped, and hypogranular platelet morphologies out of total platelets are presented. **(D)** Red blood cell morphologies including the presence of Howell-Jolly bodies (HJ Bodies), echinocytes, dacrocytes, and reticulocytes are presented as numbers counted per field. Statistical significance was analyzed using 2-way ANOVA followed by Tukey’s test for multiple comparisons: *p<0.05, **p<0.01, ***p<0.001, ****p<0.0001.

In our previous study, immunodeficient NOD.Cg-*Prkdc^scid^ Il2rg^tm1Wjl^*/SzJ (NSG) mice implanted with C1498 treated with a single course of AZA did not survive significantly longer than vehicle-treated mice indicating that a fully functional immune system may be necessary for AZA efficacy^12^. We repeated this experiment with long-term AZA treatment to ensure higher penetrance and found, again, that survival of C1498-AZA NSG mice did not differ significantly from C1498-vehicle NSG mice (**Supplementary Figure 1A**), highlighting the importance of immunocompetency for AZA efficacy (**Supplemental Figure 1A**).

Hematologic recovery of blood cell types is an important indicator of AZA efficacy and AML remission. As it takes approximately three weeks for a minimal number of actively growing C1498 blasts to cause end stage AML, we interpolated that mice should be in remission/peak AZA efficacy after two consecutive weeks of treatment. Peripheral blood was drawn 48 hours after the last vehicle or AZA treatment and WBC counts were conducted (**Figure 1B**). At this timepoint, very few C1498 blasts were seen in either C1498-vehicle or C1498-AZA mice (0.99% and 0.32%, respectively). C1498-AZA blood showed significantly decreased percentages of eosinophils (0.44% vs 3.15%) and monocytes (1.88% vs 6.17%) and an increased percentage of lymphocytes (93.8% vs 86.0%) compared to C1498-vehicle mice, respectively. However, the percentages of eosinophils (0.44% vs 0.00%) and neutrophils (3.45% vs 1.12%) were still significantly higher in C1498-AZA mice compared to healthy mice, with lymphocytes concurrently lower (93.8% vs 97.7%). These data could indicate that some AZA-treated mice do not achieve complete hematologic recovery at this timepoint. Interestingly, blood smears done at week two from C1498-AZA NSG mice were almost completely devoid of WBCs. C1498-vehicle NSG mice showed that blood was populated mostly by monocytes and neutrophils (**Supplemental Figure 1B**) as is to be expected in immunodeficient NSG mice^31^. These data may indicate that AZA treatment removes these immunosuppressive cell types but a lack of functional lymphocytes impedes the clearance of C1498s. This conclusion appears to agree with data from the immunocompetent model wherein AZA treatment, minimizes the expansion of neutrophils and maintains a higher abundance of lymphocytes compared to vehicle treatment.

Total platelets did not differ significantly between C1498-vehicle, C1498-AZA, or healthy mice but the percentage of hypogranular platelets was significantly higher in C1498-AZA mice compared to C1498-vehicle mice (**Figure 1C**). Although not significant, several C1498-AZA blood smears showed a high number of echinocytes (**Figure 1D**), possibly due to a direct effect of AZA treatment on RBCs as echinocytes can be found after exposure to other chemotherapeutic agents^32,33^. Overall, these data suggest that therapeutic doses of AZA, leading to remission in leukemic mice, re-establishes profiles resembling a healthy blood microenvironment.

### AZA Treatment Generates an Anti-Leukemic Immune Cell Profile in Blood and Decreases Overall Immune Cellularity in the Spleen

While the bone marrow microenvironment is a key player in leukemogenesis and AML progression, human datasets comprised of bone marrow mononuclear cells (typically >80% leukemic blasts) before and after HMA treatment have been analyzed for biomarkers of HMA efficacy with no significant gene expression changes found^34^ (GSE77750 AML samples only, GSE116567). The study of the blood and spleen microenvironments in AML patients have been neglected despite the presence of leukemic blasts in the peripheral blood and spleen compartments^12,35,36^, as well as the spleen’s importance as a second site for normal and/or malignant hematopoiesis^37^.

To delve more deeply into the phenotypes of WBCs and other blood components in peripheral blood and spleen, RNA isolated from whole blood or whole spleen samples from C1498-vehicle, C1498-AZA, or healthy, immunocompetent mice was assayed for gene expression using a panel of 543 immune related genes. The genes in this panel focused on the cell activation, cytokine production, adaptive immune response, immune effector processes, activation and regulation of the immune response, and leukocyte differentiation (**Supplemental Figure 2**).

Although peripheral blood cell types were enumerated by manual CBC **in Figure 1B**, deconvolution of cell types from bulk RNA reveals differences in WBC phenotypes that cannot be differentiated by eye (**Figure 2**). In blood samples, most C1498-AZA samples clustered with Healthy with regard to cell type score whereas one C1498-AZA sample clustered with C1498-DMSO samples (**Figure 2A**). When comparing specific cell type scores, C1498-AZA blood had significantly increased CD8+ T-cells, decreased Th1 cells, decreased regulatory T-cells (Treg), and decreased neutrophils compared to C1498-vehicle blood (**Figure 2B**). In spleens, Healthy and C1498-vehicle samples clustered together overall with regard to cell type scores (**Figure 2C**). C1498-AZA mice had significantly lower scores for most cells types analyzed including CD45, T, CD8+ T, Th1, cytotoxic, exhausted CD8+, Treg, B, macrophages, neutrophils, and CD56 dim NK cells (**Figure 2D**). This is not surprising, as AZA has been shown to significantly affect hematopoietic stems cells (HSCs) in the spleen, causing their differentiation and death and, ultimately, reducing overall spleen immune cellularity^38^. Overall, AZA treatment in blood appears to decrease the frequency of suppressive cell types in the blood (neutrophils and Tregs), while increasing anti-tumor cell types like CD8+ T-cells, which is conducive for anti-leukemic immunity. In the spleen, AZA decreases splenic immune cellularity, possibly through induction of differentiation and death in HSCs, a mechanism that has also been observed with leukemic stem cells (LSCs) that contributes further to its efficacy ^39^.

**Figure 2.**
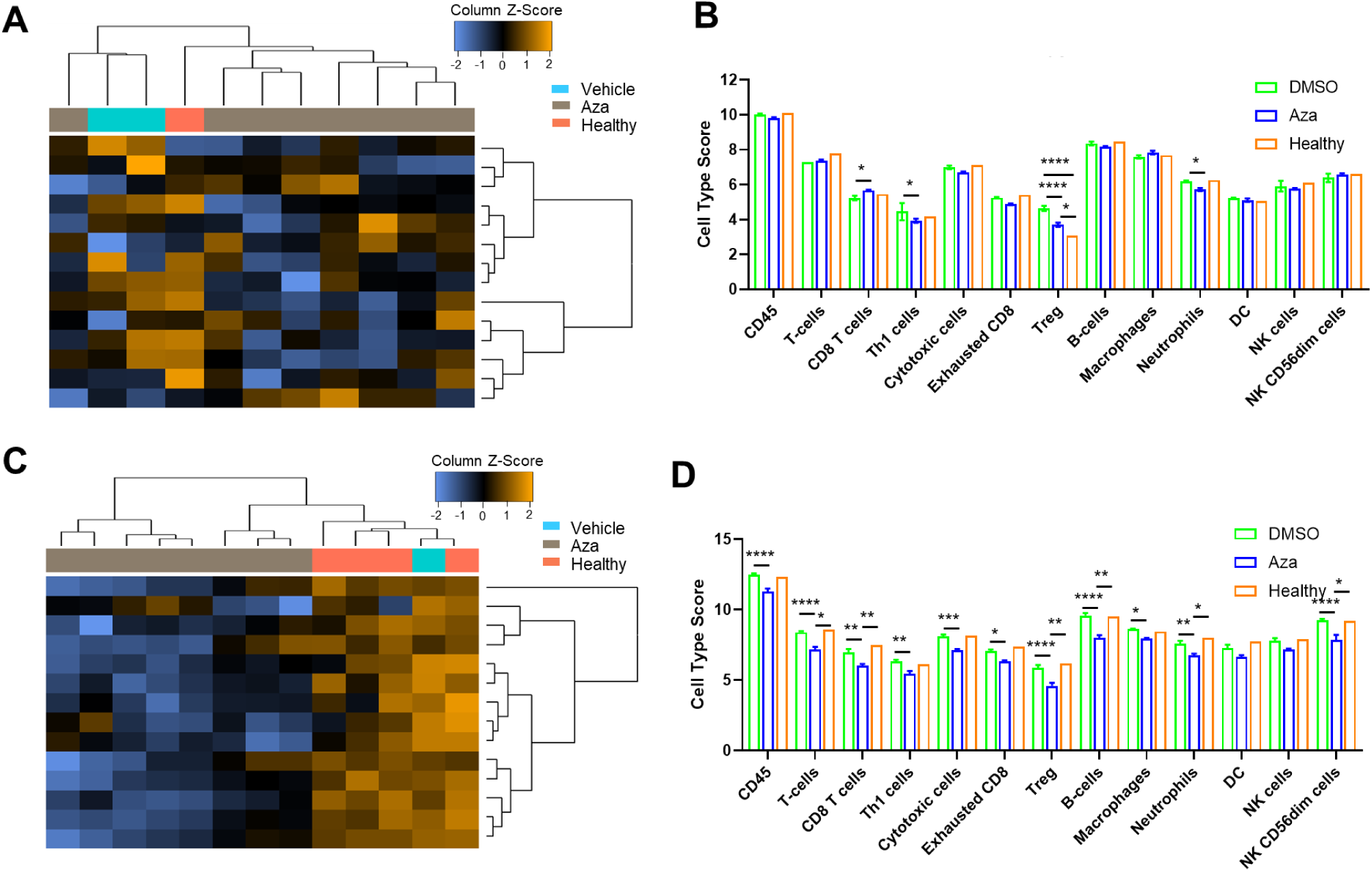
In-depth Analysis of Immune Cell Abundance and Phenotypes by Gene Expression. C1498 (80,000 cells) were implanted in C57Bl/6 mice via tail vein injection. Treatment with DMSO in diluent (Vehicle) (n=4) or 5mg/kg/mouse AZA (n=8) by intraperitoneal injection began on day 3 after implantation and continued thrice weekly. Forty-eight hours after the 6^th^ treatment (week 2) mice were euthanized after harvesting whole blood and whole spleens. Whole blood and spleen were also isolated from a non-leukemic, untreated (Healthy) mouse. mRNA was isolated from blood and spleen and probed for gene expression using the Mouse PanCancer Immune Profiling Panel. Cell type scores were calculated from bulk mRNA expression per sample and are presented as heatmaps and bar graphs for both blood **(A,B)** and spleen **(C,D)**. Statistical significance was analyzed using 2-way ANOVA followed by Tukey’s test for multiple comparisons: *p<0.05, **p<0.01, ***p<0.001, ****p<0.0001.

### Gene Pathway Alterations in Blood Induced by AZA Treatment are Largely Anti-Tumorigenic

Using gene expression data from the large-scale immune panel assay, weighted scores were developed for multiple immune response pathways (**Figure 3**). In blood, C1498-vehicle and Healthy blood samples clustered together with regard to overall pathway scores (**Figure 3A**). Compared to both C1498-vehicle and healthy blood samples, C1498-AZA mice scored significantly higher in adhesion, apoptosis, B-cell functions, cancer progression, CD molecules, cell cycle, leukocyte functions, macrophage functions, senescence, T-cell functions, TNF superfamily, and transporter functions. Only in dendritic cell and NK cell functions did C1498-AZA samples score higher than C1498-vehicle but not Healthy. Compared to C1498-vehicle and healthy samples, C1498-AZA mice scored significantly lower in adaptive immune pathways, basic cell functions, chemokines and receptors, cytokines and receptors, humoral immune pathways, inflammation, innate immune pathways, and interleukins. In the complement pathway and MHC signaling, C1498-AZA samples scored significantly lower than C1498-vehicle but not Healthy (**Figure 3B**). In spleens, almost every pathway score was significantly lower in C1498-AZA samples compared to both C1498-vehicle and Healthy samples (**Figure 3C, D**), except for basic cell functions which were higher in C1498-AZA.

**Figure 3.**
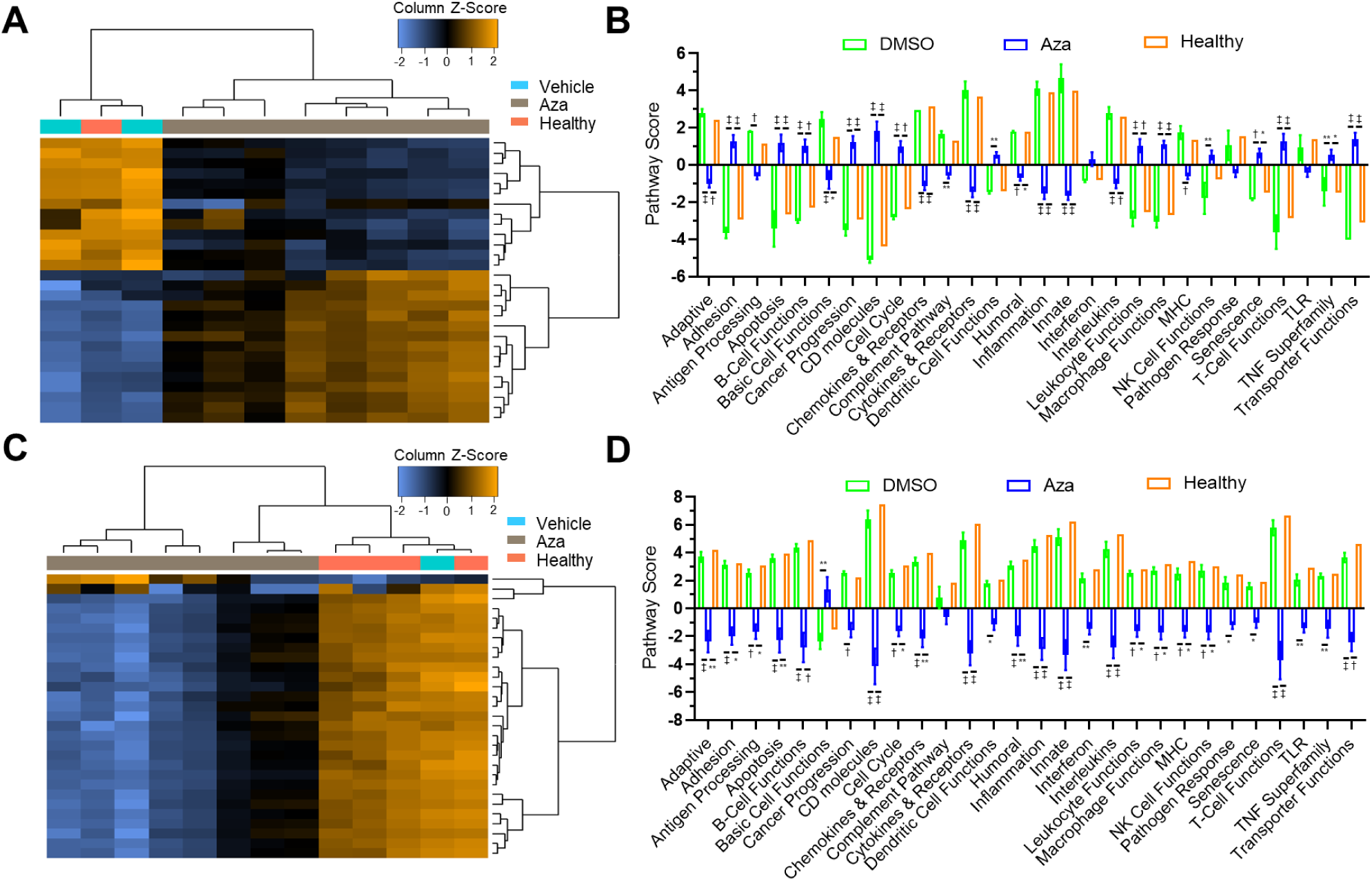
In-depth Analysis of Immune Pathways by Gene Expression. Expression levels of genes from the Mouse PanCancer Immune Profiling Panel were used to calculate scores for various immune-related pathways. Pathway scores are presented as heatmaps and bar graphs comparing non-leukemic, untreated (Healthy), DMSO-treated leukemic mice (Vehicle), and AZA-treated leukemic mice (AZA) for blood **(A,B)** and spleen **(C,D)** samples. Statistical significance was analyzed using 2-way ANOVA followed by Tukey’s test for multiple comparisons: *p<0.05, **p<0.01, †p<0.001, ‡p<0.0001.

Mirroring the decrease in pro-tumorigenic, immunosuppressive cell types in AZA-treated blood, decreased scores in inflammation and innate immune pathways along with the increase in T- and B-cell functions appears to indicate a more anti-tumor phenotype. There are conflicting data, however, such as decreased scores for adaptive immune signaling and basic cell functions, with increased scores for cancer progression and macrophage functions. These could indicate the persistence of suppressive macrophages that decrease the ability of lymphocytes to transmit signals to downstream players in adaptive immunity. A global decrease in pathway scores for C1498-AZA spleen samples also mirrors the downregulation of white blood cells seen in **Figure 2D**. Again, this is likely due to enhanced differentiation and death of HSCs induced by AZA treatment^40,41^. Taken together, these gene pathway studies highlight the complex changes occurring in the blood and spleen which may be direct consequences of AZA treatment while others may reflect compensatory mechanisms.

### Differential Gene Expression Analysis Identifies Common Targets in Blood and Spleen Modulated by AZA

When comparing specific gene expression differences in C1498-AZA compared to C1498-vehicle blood samples five candidates were significantly distinct by fold change: *Anp32b* (−2.02), *Ccr3* (−1.81), *Cfh* (1.85), *Itgb3* (2.75), and *Thbs1* (3.43) (**Figure 4A**). In spleen samples, 30 candidate genes were significantly distinct by fold change in C1498-AZA compared to C1498-vehicle: *Klra17* (−1.65), *Il5ra* (−1.64), *Cd79b* (−1.63), *Cd69* (−1.46), *Cxcr4* (−1.40), *Stat4* (−1.40), *Ms4a1* (−1.40), *Ccr2* (−1.33), *Havcr2* (−1.27), *Il16* (−1.20), *H2-Aa* (−1.17), *Bcl6* (−1.16), *Abca1* (−1.03), *Zbp1* (−1.02), *Cybb* (−0.98), *Ly9* (−0.96), *Cd244* (−0.95), *Ddx58* (−0.94), *Tnfaip3* (−0.85), *Il13ra1* (−0.81), *Ifngr1* (−0.64), *Hif1a* (−0.50), *Psmd7* (0.84), *Anp32b* (1.08), *Ccl17* (1.80), *Blm* (1.83), *Thbs1* (1.93), *Birc5* (2.06), *Tal1* (3.03), and *Mpo* (3.78) (**Figure 4B**). Two targets, *Thbs1* and *Anp32b*, are shared between the blood and spleen datasets (**Figure 4C**), and while *Anp32b* is downregulated in blood but upregulated in spleen after AZA treatment, *Thbs1* is upregulated in both blood and spleen after AZA treatment. The upregulation of *Thbs1* in blood serum has already been associated with better prognosis in AML patients undergoing AZA treatment^42^, likely due to its potent inhibition of angiogenesis^43,44^.

**Figure 4.**
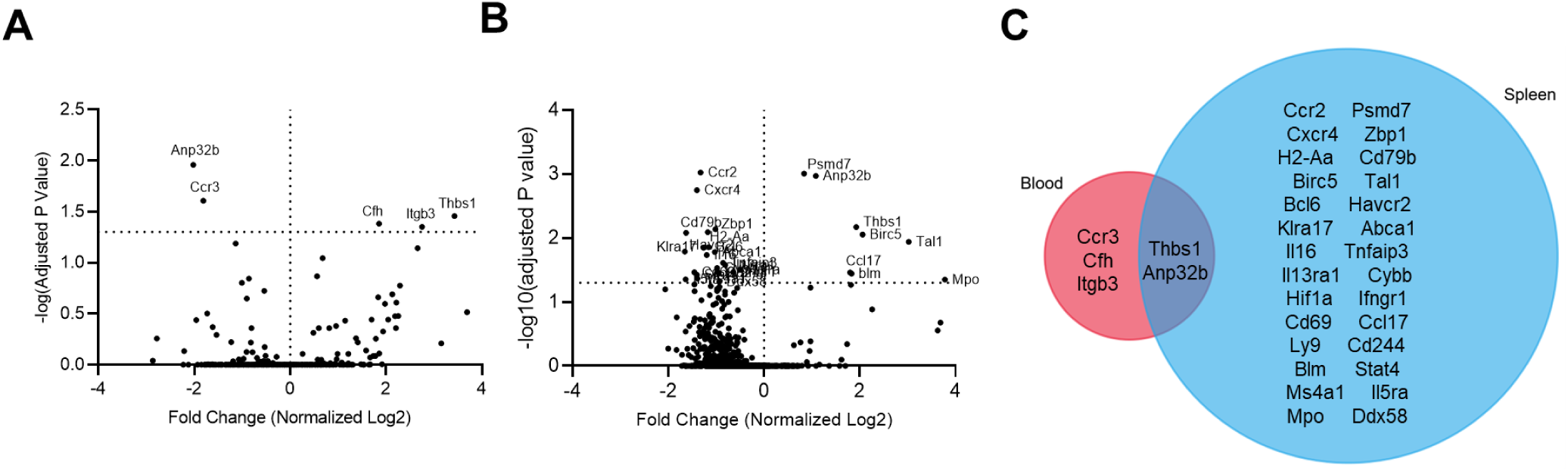
Differential Gene Expression Analysis Between Vehicle- and AZA-treated Blood and Spleen Samples from Leukemic Mice. Differentially expressed genes from the Mouse PanCancer Immune Profiling Panel between DMSO-treated leukemic mice (Vehicle) and AZA-treated leukemic mice (AZA) are shown in volcano plots as fold change (log2) versus adjusted p-values (−log10). Fold changes are presented as AZA versus vehicle for blood **(A)** and spleen **(B)** samples with significant genes labeled. Statistical significance was analyzed with multiple unpaired t-tests and corrected for multiple comparisons using the Holm-Sidak method. **(C)** A Venn diagram of genes shared between blood and spleen samples is presented.

### Gene Targets Modulated by AZA are Primarily Associated with Cell Surface Adhesion and Signaling, Thrombosis, and Angiogenesis

A functional enrichment analysis of AZA-targeted genes in blood revealed that their gene products are expressed on the surfaces of cells and in platelets (from the gene subontology “Cellular Component”), and are involved in fibrinogen, fibronectin, cytokine, fibroblast growth factor, and extracellular matrix binding (from the gene subontology “Molecular Function”). These functions are implicated in the control of vasculature development and angiogenesis, including endothelial cell proliferation and apoptosis, as well as fibroblast migration, lipid transport, apoptotic cell clearance, blood coagulation and platelet aggregation, and chemotaxis (form the gene subontology “Biological Processes”) (**Figure 5A**).

**Figure 5.**
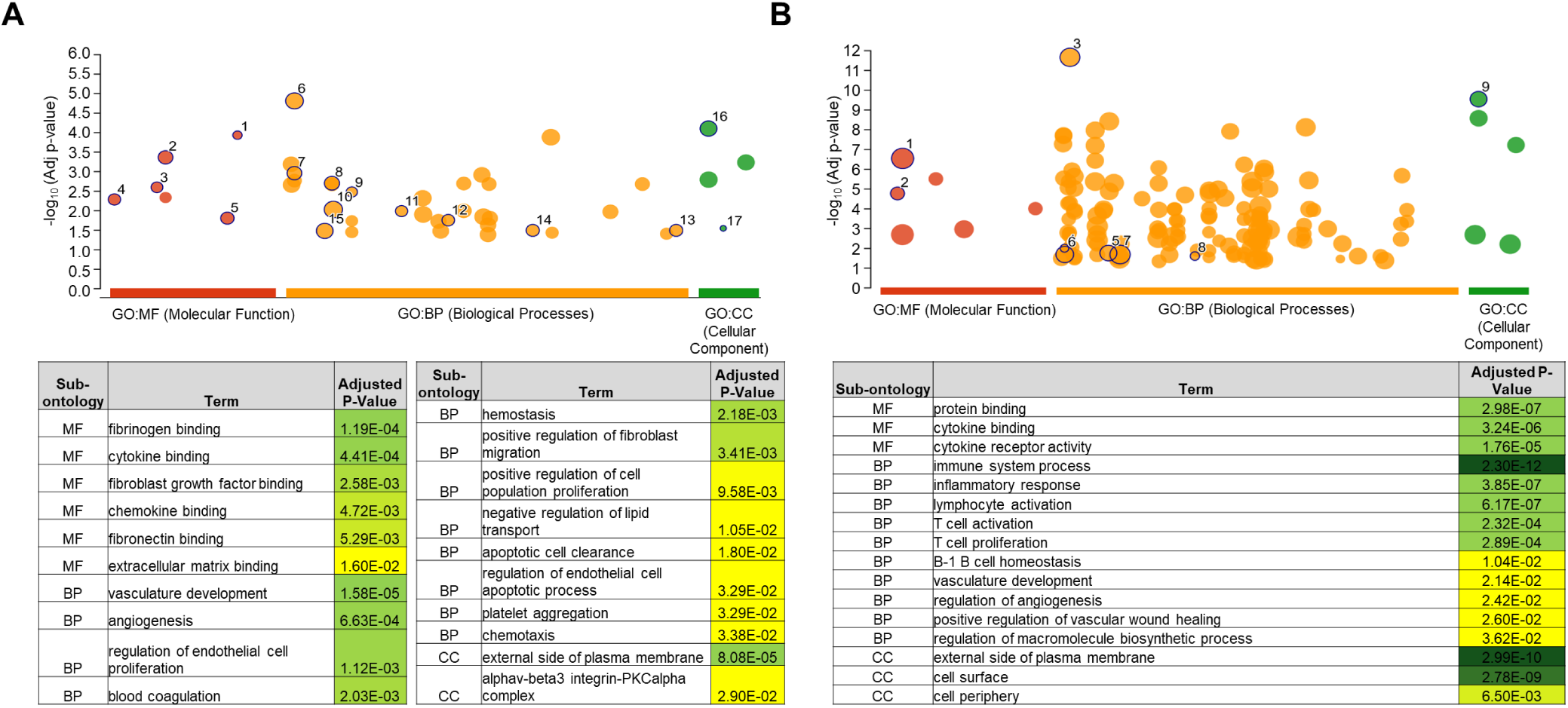
Functional Enrichment Analysis of Genes Differentially Expressed Between Vehicle- and AZA-treated Blood and Spleen Samples from Leukemic Mice. Differentially expressed genes from the Mouse PanCancer Immune Profiling Panel between DMSO-treated leukemic mice (Vehicle) and AZA-treated leukemic mice (AZA) were entered into the g:GOSt web tool for gene set enrichment analysis. Statistically significant enriched Gene Ontology (GO) terms for blood **(A)** and spleen **(B)** are presented as a pictogram highlighting all enriched terms, and a table of driver terms with adjusted p-values. Statistical significance of enrichment was determined using the default g:SCS algorithm.

AZA-targeted genes in spleen were enriched for expression on cell surfaces and function in protein binding and cytokine receptor activity. The biological processes that they are enriched for include general immune system processes, B-1 B cell homeostasis, response to xenobiotic stimulus, vasculature development, positive regulation of macromolecule metabolic processes, and positive regulation of vasculature wound healing (**Figure 5B**).

CFH and THBS1, despite being positive regulators of platelet aggregation^45,46^, both strongly inhibit angiogenesis^47,48^. ANP32b, called APRIL in humans, is thought to be a promoter of angiogenesis^49^. Downregulation of *Anp32b* and upregulation *Cfh* and *Thbs1* in C1498-AZA mouse blood could result in decreased angiogenesis. In the spleen, downregulation of the pro-angiogenic genes *Cxcr4*^50^, *Stat4*^51^, *Ccr2*^52^, *Bcl6*^53^, *Hif1a*^54^ and *Il16*^55^, as well as upregulation of anti-angiogenic *Thbs1* supports that the mechanism for AZA efficacy may involve inhibiting angiogenesis.

### Gene Modulation by AZA Alters Leukemic Disease Outcome Following Relapse

To determine how AZA-driven gene expression modulation and possible inhibition of angiogenesis affects disease progression, blood and target tissues from C1498-Vehicle and C1498-AZA mice were examined. At terminal stages of disease, C1498-AZA and C1498-vehicle mice show no significant difference in expression of the target genes in whole blood samples (**Supplemental Figure 3**). Indeed, amounts of C1498 cells found in terminal blood draws did not differ between C1498-vehicle and C1498-AZA mice, and make up about 6.63% of total WBCs (**Figure 6A**). However, even at end stage disease, AZA treatment limited the expansion of neutrophils while maintaining a higher percentage of lymphocytes compared to C1498-vehicle (13.4% vs 28.4% and 73.6% vs 53.2%, respectively), although not to the level of healthy mice. Total platelet counts did not differ between C1498-vehicle, −AZA, or healthy mice, but platelet morphology did differ significantly with both C1498-vehicle and C1498-AZA mice showing significantly higher percentages of hypogranular platelets with concurrent lower percentages of normal platelets compared to healthy (**Figure 6B**). RBCs in C1498-AZA mice showed a significant increase in echinocyte morphology compared to both C1498-vehicle and healthy blood smears (**Figure 6C**).

**Figure 6.**
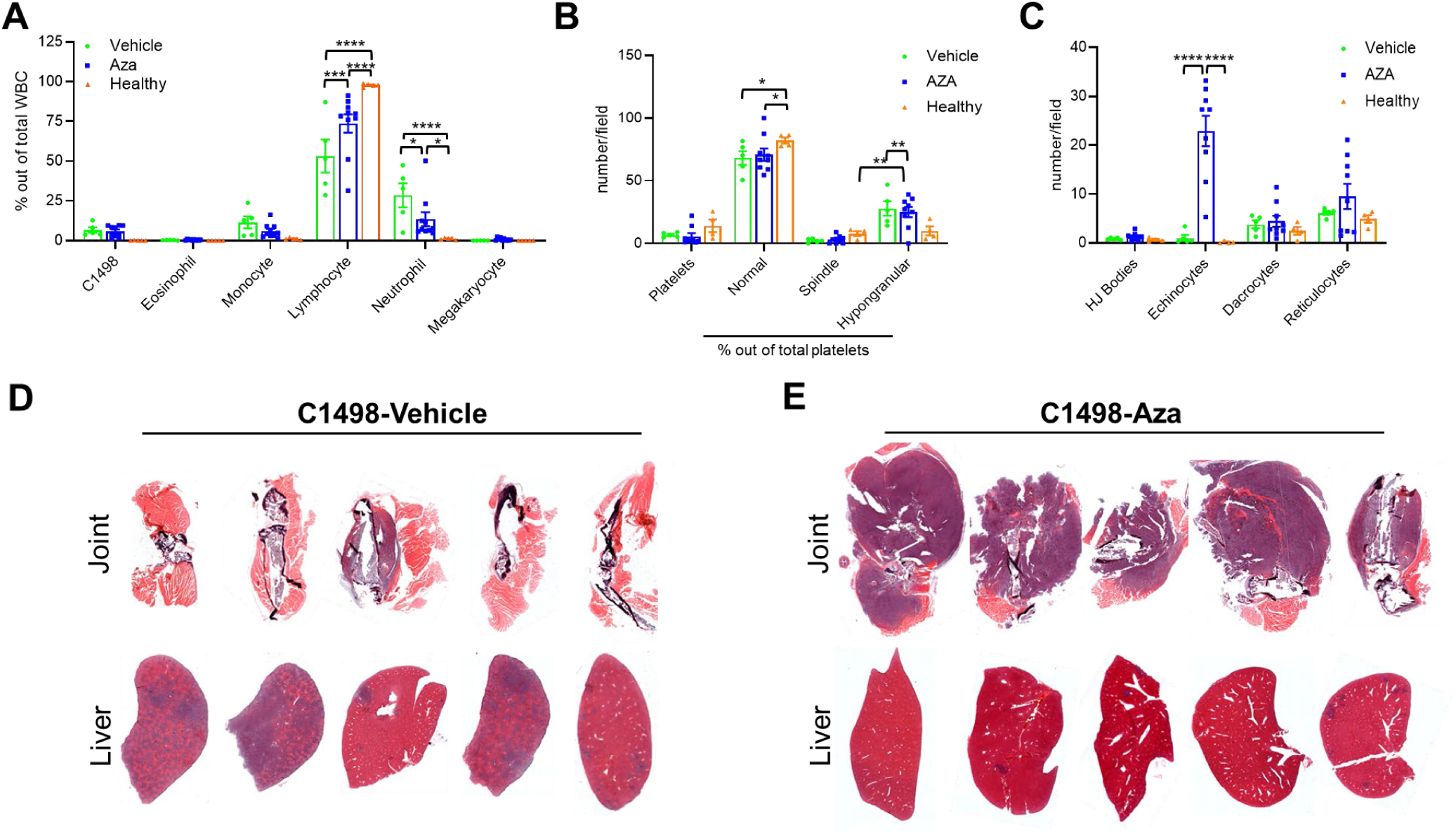
Hematologic Profiles and Tissues from Terminally Leukemic Vehicle- and AZA-treated mice. C1498 cells (80,000) were implanted in C57Bl/6 mice via tail vein injection. Treatment with DMSO in diluent (Vehicle) (n=5) or 5mg/kg/mouse AZA (n=11) by intraperitoneal injection began on day 3 after implantation and continued thrice weekly until survival endpoints. Mice were euthanized once they showed signs of terminal disease including, but not limited to, labored breathing, a hunched posture, or decreased mobility. Whole blood was taken just prior to euthanasia, smeared onto slides and stained for complete blood counts. **(A)** White blood cells (WBC) were counted manually, including C1498, and are presented as a percentage of total WBC counted. **(B)** Numbers of platelets and percentages of normal, spindle-shaped, and hypogranular platelet morphologies out of total platelets are presented. **(C)** Red blood cell morphologies including the presence of Howell-Jolly bodies (HJ Bodies), echinocytes, dacrocytes, and reticulocytes are presented as numbers counted per field. Statistical significance was analyzed using 2-way ANOVA followed by Tukey’s test for multiple comparisons: *p<0.05, **p<0.01, ***p<0.001, ****p<0.0001. Multiple organs and tissues including hip and knee joints and livers were removed post-euthanasia, fixed, and stained by Hematoxylin and Eosin. Representative joint and liver tissues for **(D)** C1498-Vehicle and **(E)** C1498-AZA are shown (n=5).

Terminally ill C1498-vehicle mice displayed a hunched position, labored breathing, and lower body temperature prior to euthanasia. Liver enlargement and splenomegaly was observed in all C1498-vehicle mice during gross anatomy. Histochemical staining of organs revealed infiltration by cancerous cells with atypical nuclei, dense chromatin, and a high nuclear/cytoplasmic ratio most notably in the liver (**Figure 6D**). Similar infiltrations were observed in C1498-AZA mice but to a somewhat lesser extent (**Figure 6E**). Terminally ill C1498-AZA mice did not outwardly show signs of morbidity but had to be euthanized due to mobility issues. Masses were palpable on hip and knee joints (typically a single mass per mouse), sometimes causing paralysis of the affected limb. Histologically, these masses were revealed to be cancerous cells, possibly having invaded directly from bone marrow through damaged bone, rather than metastasizing to distal tissues via the blood stream. C1498-AZA mice showed more growths of cancerous cells around bone compared to liver, with the opposite being true of C1498-vehicle mice (**Figure 6D, E**). AML can metastasize to other organs and tissues with these cases having a very poor prognosis^56,57^. As angiogenesis is tightly linked to the metastasis of AML^58^, these results could indicate that AZA decreases metastasis (to the liver in this model) via the gene expression changes that may be inhibiting angiogenesis.

### Gene Alterations after AZA Treatment in PBMC from Human AML Patients Confirm Upregulation of Thrombosis with Downregulation of Angiogenesis Pathways

Despite most available human datasets containing only leukemic burdened bone marrow or sorted leukemic blasts, we did find one dataset of PBMCs harvested from AZA responsive AML patients. Evaluating RNAseq expression data from these patients after three complete cycles of AZA treatment compared to pre-treatment, we identified a list of genes that were significantly changed (**Figure 7A**, **Supplemental Table 1**). Out of the 340 genes found to be significantly affected by AZA treatment, only *ITGB3* was a common gene upregulated in our AZA-targeted gene candidates in blood and the patient gene set (**Figure 7B**), and only the *STAT4* gene was common between our AZA-targeted spleen gene candidates and the human gene set (**Figure 7C**). Interestingly, the expression of both *ITGB3* and *STAT4* gene products from non-malignant cell types in the tumor microenvironment has previously been shown to support anti-tumor immunity^59–61^. When genes from the human dataset were analyzed for functional enrichment, overlap was observed in the subontologies of Molecular Function and Cellular Component (**Figure 7D**) with both the blood and spleen data sets from mice, namely enrichment in cytokine and chemokine binding and activity and expression on the cell periphery and in platelet alpha granules. A gene set enrichment analysis (GSEA) showed that several pathways were upregulated in AZA-treated PBMC compared to pre-treated (**Figure 7E**). including T-cell activation, inflammation mediated by chemokines and cytokines, the Alzheimer disease-presenilin pathway, blood coagulation, and platelet-derived growth factor signaling. Downregulated pathways included apoptosis signaling, integrin signaling, heterotrimeric G-protein signaling pathways mediated by Gi alpha/Gs alpha and Gq alpha/Go alpha, and angiogenesis. These data, especially upregulation of T-cell activation, cytokine and chemokine signaling, and blood coagulation concurrent with downregulation of angiogenesis agree with our findings using the C1498 AML model.

**Figure 7.**
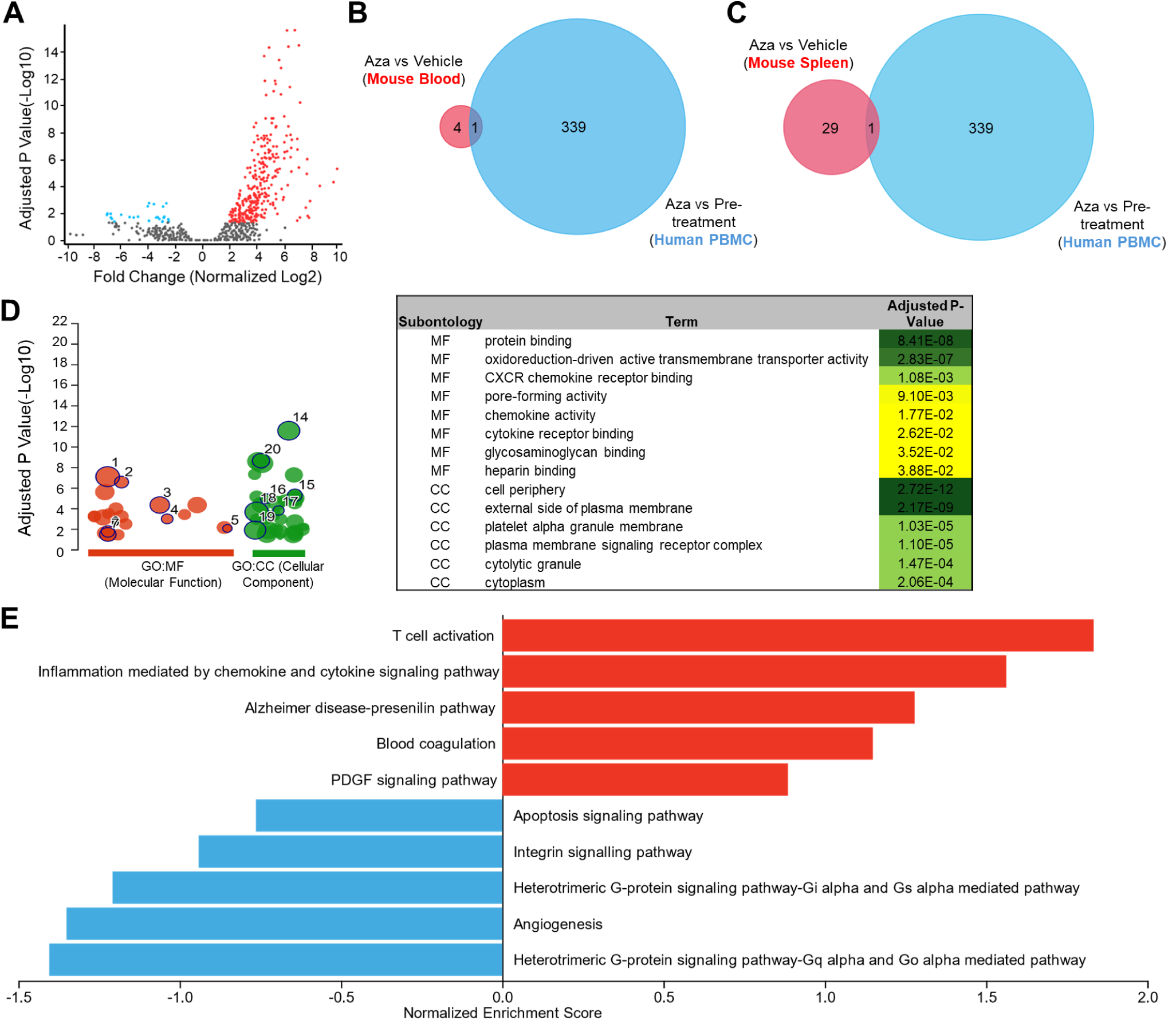
Analysis of Genes Altered by AZA Treatment in Human AML. Gene Expression Omnibus (GEO) data set GSE118558 was analyzed for significant differences in gene expression comparing peripheral blood mononuclear cells (PBMC) from AZA-responsive AML and MDS patients after three complete cycles of AZA treatment versus pre-treatment. **(A)** Volcano plot of normalized gene expression (Log2) against adjusted p-values (−Log10) for AZA treated versus pre-treatment PBMCs. Significantly upregulated genes are highlighted in red, significantly downregulated genes are highlighted in blue, non-significant genes are shown in grey. P-values were adjusted using the Benjamini and Hochberg method for false discovery rate with a cut-off of 0.05. **(B)** Venn diagram of significant differentially expressed genes from the mouse blood dataset comparing AZA- vs DMSO-treated leukemic mice and the human PBMC dataset comparing AZA-treated to pre-treated AML and MDS patients. **(C)** Venn diagram of significant differentially expressed genes from the mouse spleen dataset comparing AZA- vs DMSO-treated leukemic mice and the human PBMC dataset comparing AZA-treated to pre-treated AML and MDS patients. **(D)** The 340 genes differentially expressed in AML and MDS patients responsive to AZA treatment in PBMC post-three cycles of treatment versus pre-treatment, were entered into the g:GOSt web tool for gene set enrichment analysis. Statistically significant enriched Gene Ontology (GO) terms for are presented as a pictogram highlighting all enriched terms, and a table of notable terms with adjusted p-values. Statistical significance of enrichment was determined using the default g:SCS algorithm. **(E)** Significantly affected genes from the human PBMC dataset were subjected to Gene Set Enrichment Analysis. Top pathways are graphed against enrichment scores to be interpreted as up- or down-regulated pathways.

## DISCUSSION

Gene alterations driving AML leukemogenesis and progression frequently dysregulate histone-modifiers and other epigenetic regulatory proteins (including at the level of promoter enhancers)^62–67^, this leads to unique variations on gene expression between AML clones within and between patients^68,69^. Hence, the prognostic value of specific or global gene methylation or expression patterns after HMA treatment has been limited ^70,71^.

In this study we found that analysis of the leukemia microenvironment in whole blood and spleen revealed altered expression of genes related to cytokine and chemokine binding, cell adhesion, thrombosis and angiogenesis in non-leukemic blood cells and splenocytes after AZA treatment. Literature on individual genes reveals their involvement as pro- or anti-thrombosis and/or angiogenesis, overall seeming to indicate an upregulation or maintenance of normal thrombotic processes (usually deficient in AML patients^72^) with an inhibition in angiogenesis. Analysis of human genes affected by AZA treatment in blood, although specifying a largely different set of genes, confirms an upregulation in blood coagulation pathways with a downregulation in angiogenesis.

Further studies are needed to characterize remaining immune cells, platelets, and their association with endothelial cells and angiogenesis after AZA treatment, but overall it appears that a major caveat of the AZA mechanism of action for prolonging survival in AML cases involves decreasing angiogenesis while maintaining healthy platelet function.

Interestingly, human AML blasts express *Cfh*, *Ccr3*, and *Thbs1*, and higher expression of all are separately associated with poor prognosis, including *Stat4* ^51,73–75^. Expression of *Anp32b* is increased in AML blasts and higher expression correlates with poorer overall survival^76^, and expression of *Ccr3* in AML blasts is thought to drive chemokine induced proliferation^77,78^. Thus, systemic targeting of these genes or their pathways should have effects on both leukemic cells and the normal cell types utilizing these gene products for anti-leukemic processes. Great care should be taken to fully understand the interplay of adhesion, thrombosis, and angiogenesis between leukemic cells, normal immune cells, and other stromal cells before rational combination therapies with AZA can be safely explored in the clinic.

## METHODS AND MATERIALS

### Animals and cell lines

C57Bl/6 mice were obtained from breeding colonies at the City of Hope (COH) Animal Research Center. NOD/SCID gamma (NSG) mice were obtained from Jackson Labs (catalog# 005557). All experimental mice were 6-8 weeks old. Mice were handled according to standard institutional animal care and use committee (IACUC) guidelines (protocol #17128). The C1498 cell line was obtained from ATCC® (TIB-49, Manassas, VA, USA). FBL3 cells were obtained from the DCTD tumor repository (NCI Frederick, MD, USA). Both cell lines were maintained in RPMI media supplemented with 10% FBS, 2mM L-glutamine and 100 units/mL penicillin, and 100 ug/mL streptomycin. Cell lines were passaged minimally (≤5 times) before implantation in mice.

### In vivo 5-Aza treatment studies and survival studies following leukemia challenge

For therapeutic studies, mice were implanted with 8×10^4^ C1498 or 1×10^4^ FBL3 cells via intravenous (i.v.) tail vein injection. Three days post-implant, mice were treated with 5 mg/kg 5-Aza or Vehicle (DMSO in diluent) for 3 consecutive days by intraperitoneal (i.p.) injection in a total volume of 500µl in 1x HBSS. For survival analysis, mice showing signs of terminal disease states following leukemic challenge (lethargy, paralysis, hunched posture, etc. or ovarian mass (females)) were immediately euthanized. Survival data is displayed as Kaplan-Meier curves.

#### Complete Blood Counts

Whole blood was sampled from the retroorbital vein using heparinized microhematocrit capillary tubes (22-362-566, Fisherbrand; Waltham, MA, USA). Ten microliters were smeared onto lysine-coated glass slides. The blood smear was allowed to dry and the slide was fixed in methanol followed by staining with Wright-Giemsa (WG32, Sigma-Aldrich; St. Loius, MO, USA). Complete blood counts were conducted manually using a light microscope. Platelet counts and morphology observations were made from images taken on the Widefield Observer 7 (Zeiss; Oberkochen, Germany) microscope using the 100x lens with oil. The AxioCam 506 Color camera (Zeiss) was used to capture images. Images were analyzed using ZEN Blue 3.0 software (Zeiss).

### Quantitative PCR

Total RNA from whole blood was isolated using the Blood RNA Mini Kit (R02-01, Bioland Scientific; Paramount, CA, USA). Total RNA was converted to cDNA using the High Capacity cDNA Reverse Transcription Kit (using random primers) (4368814, Applied Biosystems; Waltham, MA, USA). cDNA was amplified and fluorescently labeled using the Power-Up™ 2x SYBR™ Green Mastermix for qPCR (containing ROX) (A25742, Applied Biosystems; Waltham, MA, USA). Experiments were run with “no template” controls to control for nucleic acid contamination in water or primers, no data was used if sample CTs fell within five CTs of the “no template” controls. Primer sequences: (1) *Anp32b* (NM_130889.3) Forward - TTAAGCGCAGTGCGGGG Reverse – AAGACAAGTTCTCGAACGGCTG; (2) *Ccr3* (NM_009914.4) Forward – ACGGAAGAACTGTAACAAGTTGG Reverse – GCCATTCTACTTGTCTCTCTGGTGA; (3) *Cfh* (NM_009888.3) Forward – GGATCCACCACATGTGCCAA Reverse – ATTTCCCTGTTGAGTCTCGGC; (4) *Itgb3* (NM_016780.2) Forward - CAGTGGCCGGGACAACTC Reverse – ATCTTCGAATCATCTGGCCGT; (5) *Thbs1* (NM_001313914.1) Forward – CAGCATCCGAAAAGTGACGG Reverse – GGCAGGTCCTTGTCTGTACC.

### Statistics

All statistical analyses were performed using the Prism software by GraphPad (V9) (San Deigo, CA, USA) unless otherwise specified in figure legends. Unless otherwise indicated, all error bars represent standard error of the mean.

### Network Topology Analysis of Mouse PanCancer Immune Profiling Panel

The full gene list, unranked, included in the Mouse PanCancer Immune Profiling Panel was entered into the WEB-based Gene Set Analysis Toolkit (WebGestalt.org)^79^ for Network-Topology-based Analysis. Mus musculus was selected as the organism of interest and PPI BIOGRID was selected as the functional database.

### Bulk RNA Gene Expression Analysis for Mouse Blood and Spleen

RNA Extraction and QC: The RNeasy Plus Mini kit (74134, Qiagen; Hilden, Germany) was used to extract RNA following the manual of manufacture for mouse samples of whole blood or spleen. The extracted RNA was quantified using the Qubit RNA high-sensitivity kit (Q32852, ThermoFisher; Waltham, MA, USA) and the Qubit 4 Fluorometer (ThermoFisher). Size distribution was measured using Bioanalyzer 2100 (Agilent; Santa Clara, CA, USA).

nCounter Bulk RNA Expression Assay: The CodeSet hybridization was set up by mixing 8ul of total RNA (200ng); 10ul hybridization master mix containing 5ul of hybridization buffer, 3ul of Reporter CodeSet and 2ul of Reporter Plus CodeSet; and capture master mix (100052, Nanostring; Seattle, WA, USA) containing 2ul of Capture Probe Set and 1ul of Capture Plus Probe Set. The CodeSet of Mouse PanCancer Immune Profiling Panel with Panel Plus (20 additional genes, Supplemental Table 2) was used for this study (15000142, Nanostring). After overnight hybridization at 65C for 18h, the hybridized samples were processed in the prep station and the data acquisition was performed by nCounter MAX/FLEX profiler (Nanostring).

### Differential Gene Expression, Cell Type Score, and Pathway Score Analysis of Mouse Gene Expression Data

nCounter bulk RNA expression data were analyzed using nSolver Analysis Software (Version 4.0, R Version 3.3.2) with the nSolver Advanced Analysis Add-On (Nanostring).

### Functional Enrichment Analysis Using the g:GOst Webtool

Unranked, unordered gene lists were uploaded to https://biit.cs.ut.ee/gprofiler/gost^80^. For mouse genes sets, *Mus musculus* was chosen as the organism of interest, *Homo sapiens* was chosen for human gene sets. Queries were run for annotated genes only using Fisher’s one-tailed t-test measuring the randomness of intersection between the query and the term. P-values were adjusted for multiple hypothesis testing using the g:SCS method and a threshold of 0.05.

### Differential Gene Expression Analysis of Human Data

The Gene Expression Omnibus (GEO) dataset GSE118558^81^ was analyzed using GEO2R^82,83^ (National Center for Biotechnology Information (NCBI), a division of the National Library of Medicine (NLM) at the National Institutes of Health (NIH), Bethesda, MA, USA). Samples GSM3333205 and GSM3333197 were used for AZA-treatment responsive samples compared to GSM3333202 and GSM3333194 pre-treatment samples. GEO2R uses the Wald test followed by adjustment for multiple hypothesis testing using the Benjamini-Hochberg method (alpha cut-off 0.05).

## ACKNOWLEDGEMENTS

This research was supported by the National Cancer Institute (NCI) of the National In-stitutes of Health (NIH) under grant numbers R01CA266472, R01CA272732, R21CA256593, and R21CA293969. Research reported in this publication included work performed in the Molecular Pathology, Animal Resource Center, and Light Microscopy Digital Imaging cores supported by the NCI of the NIH under grant number P30CA033572. The content is solely the responsibility of the authors and does not necessarily represent the official views of the National Institutes of Health. The animal study protocol was approved by the Institu-tional Animal Care and Use Committee (IACUC) (IACUC# 17128, approved 4 March 2024).

