## Supplemental Figures and Tables for "Effects of Hypomethylating Agents on Gene Modulation in the Leukemic Microenvironment and Disease Trajectory in a Mouse Model of AML"

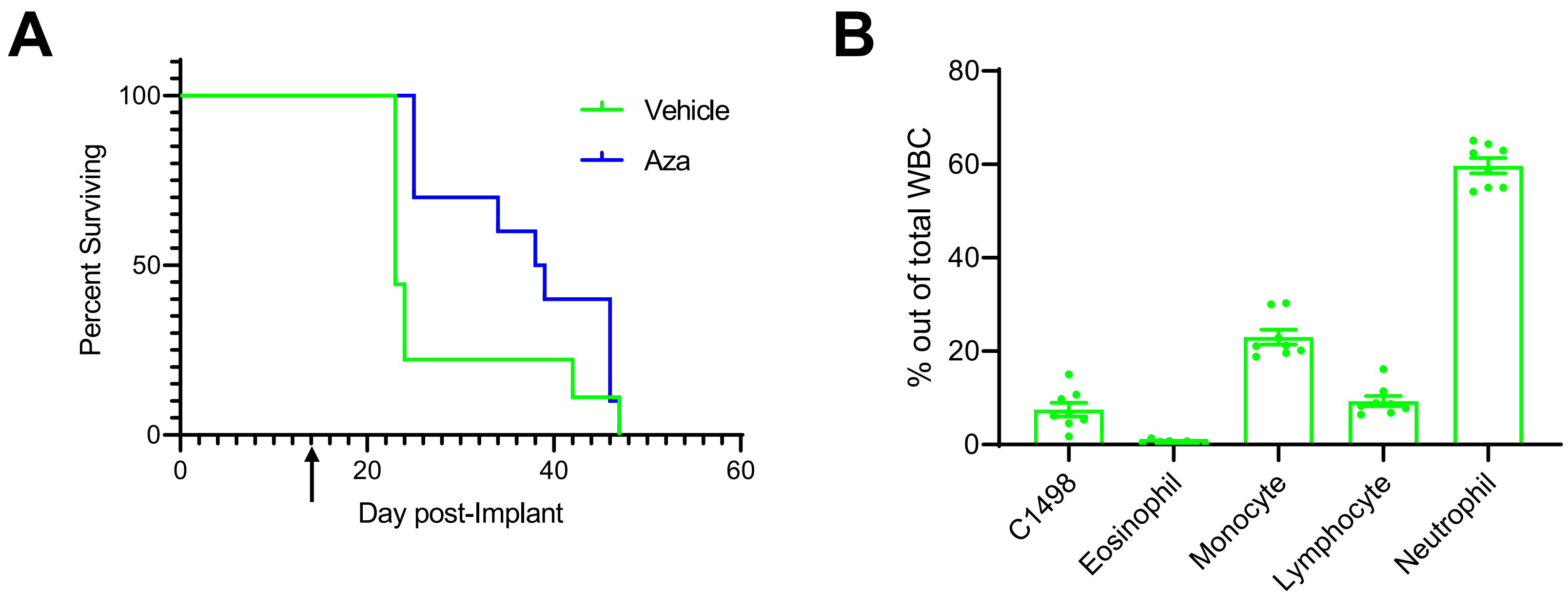


**Supplemental Figure 1. Survival and Hematologic Profiles of Leukemic Vehicle- and AZA-treated Immunodeficient Mice.** C1498 leukemia cells (80,000) were implanted in NSG immunodeficient mice via tail vein injection. Treatment with DMSO in diluent (Vehicle) (n=9) or 5mg/kg/mouse AZA (n=10) by intraperitoneal injection began on day 3 after implantation and continued thrice weekly until survival endpoints. **(A)** Mice were euthanized once they showed signs of terminal disease state and survival was recorded for each mouse and displayed as a Kaplan-Meier curve. Statistical significance was analyzed using the Log-rank (Mantel-Cox) test. The arrow (↑) indicates the timepoint at which retroorbital blood was drawn to assess hematologic profiles. **(B)** Drawn blood was stained and smeared for complete blood counts. Manually counted WBCs for C1498-Vehicle mice are shown as percentages of total WBC counted. Blood from C1498-AZA mice showed an extremely low WBC count that could not accurately be displayed graphically.

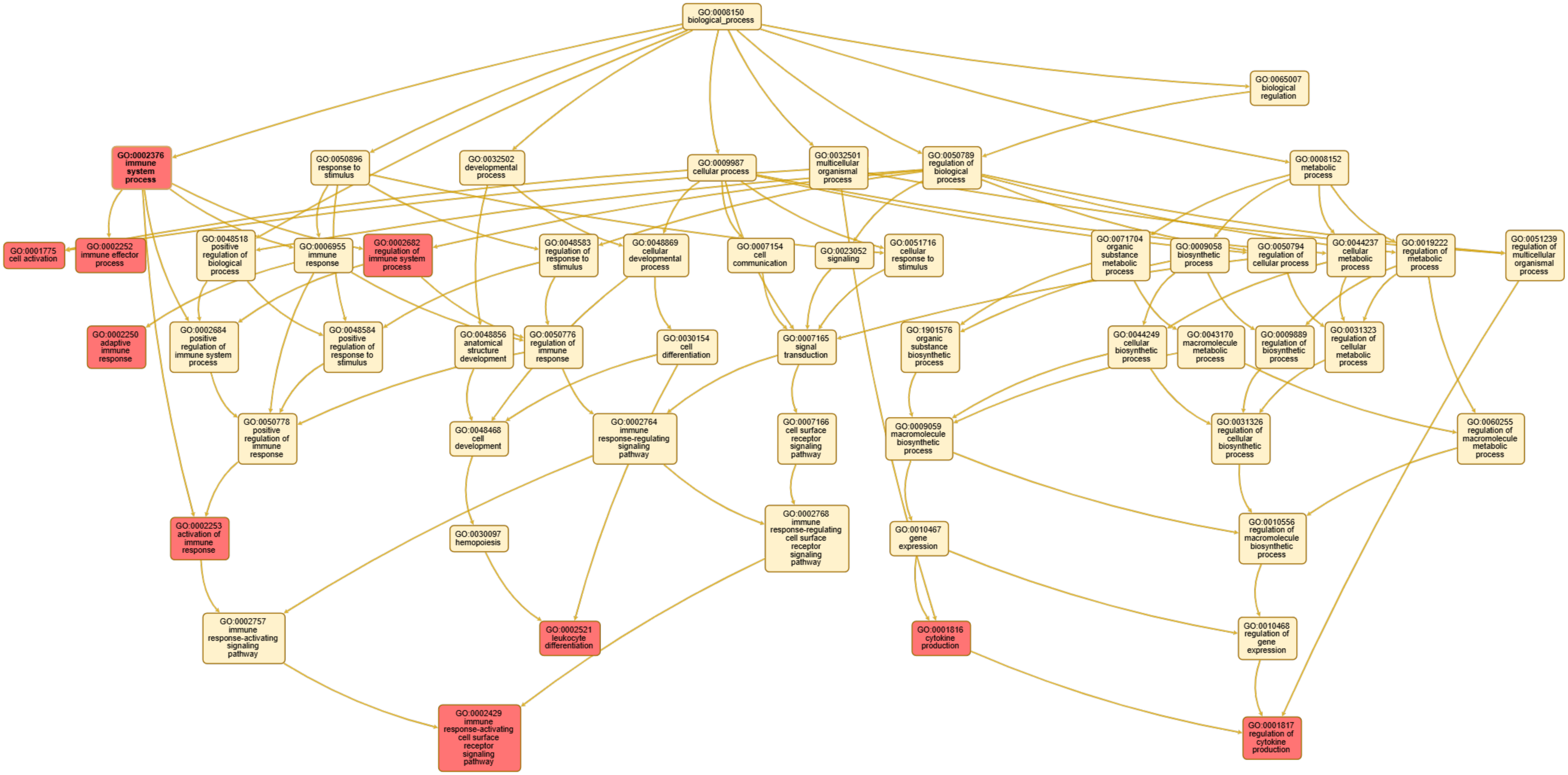


### Enriched Gene Ontology Categories

| GO Identification | GO Name |
|---|---|
| GO:0001775 | Cell Activation |
| GO:0001816 | Cytokine Production |
| GO:0001817 | Regulation of Cytokine Production |
| GO:0002250 | Adaptive Immune Response |
| GO: 0002252 | Immune Effector Process |

| GO Identification | GO Name |
|---|---|
| GO:0002253 | Activation if Immune Response |
| GO:0002376 | Immune System Process |
| GO:0002429 | Immune Response-Activating Cell Surface Receptor Signaling Pathway |
| GO:0002521 | Leukocyte Differentiation |
| GO:0002682 | Regulation of Immune System Process |

**Supplemental Figure 2. Map and Listing of Gene Ontologies Enriched in the Mouse PanCancer Immune Profiling Panel.** Genes to be probed for expression were analyzed for Network Topology using the WEB-based Gene Set Analysis Toolkit (WebGestalt). The mapping of enriched gene ontologies is shown in red with the linked ancestor ontologies in off-white. The top ten enriched ontologies are shown with their Gene Ontology (GO) identification number.

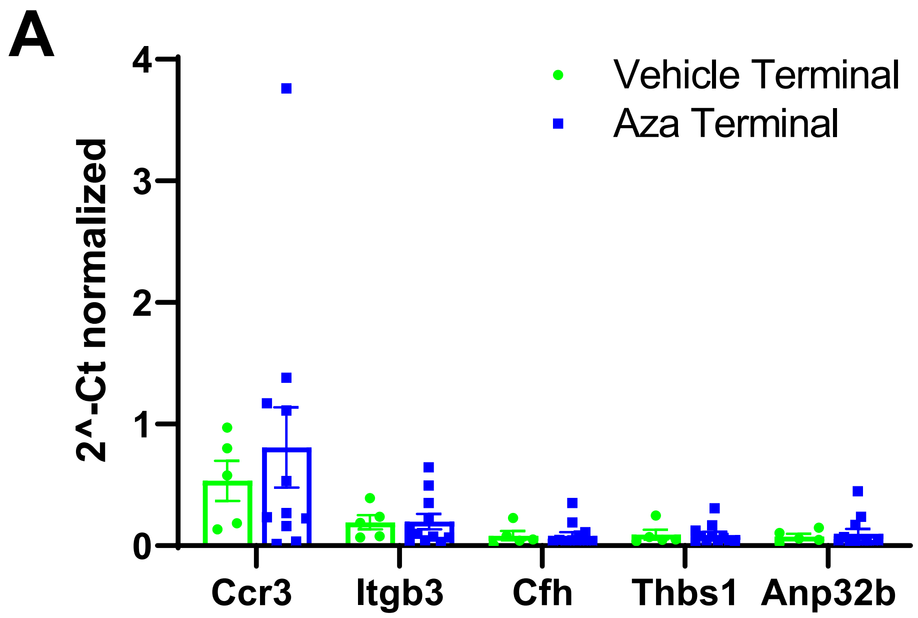


**Supplemental Figure 3. Expression of Significant Genes in Blood of Terminally Ill Mice.** C1498 cells (80,000) were implanted in C57Bl/6 mice via tail vein injection. Treatment with DMSO in diluent (Vehicle) (n=5) or 5mg/kg/mouse AZA (n=11) by intraperitoneal injection began on day 3 after implantation and continued thrice weekly until survival endpoints. Mice were euthanized once they showed signs of terminal disease including, but not limited to, labored breathing, a hunched posture, or decreased mobility. Whole blood was taken just prior to euthanasia and RNA isolated from the blood was analyzed for expression of the genes found to be significantly different between Vehicle and Aza-treated samples in the analysis of blood from week2 of treatment: *Ccr3, Cfh, Itgb3, Thbs1, Anp32b*. Expression was determined by qPCR and transformed Ct's (2^-Ct) normalized to *gapdh* are presented as a bar graph.

**Supplemental Table 1. Statistically significant genes in PBMC of human AML or MDS patients of AZA-treated samples compared to PBMCs pre-treatment.**

| GeneID | padj | pvalue | lfcSE | stat | log2Fold Change | baseMean | Symbol | Description |
|---|---|---|---|---|---|---|---|---|
| 5196 | 2.88e-16 | 2.09e-20 | 0.73 | 9.26 | 6.7220 | 914.99 | PF4 | platelet factor 4 |
| 81027 | 2.99e-16 | 4.35e-20 | 0.67 | 9.18 | 6.1426 | 954.86 | TUBB1 | tubulin beta 1 class VI |
| 5473 | 3.79e-15 | 8.26e-19 | 0.79 | 8.86 | 7.0157 | 5007.51 | PPBP | pro-platelet basic protein |
| 10398 | 4.75e-15 | 1.38e-18 | 0.71 | 8.80 | 6.2499 | 225.77 | MYL9 | myosin light chain 9 |
| 4603 | 5.53e-15 | 2.01e-18 | 0.55 | 8.76 | 4.8364 | 403.83 | MYBL1 | MYB proto-oncogene like 1 |
| 4512 | 2.35e-14 | 1.02e-17 | 0.52 | 8.57 | 4.4742 | 880614.15 | COX1 | cytochrome c oxidase subunit I |
| 64919 | 4.61e-14 | 2.34e-17 | 0.66 | 8.48 | 5.5624 | 301.09 | BCL11B | BCL11 transcription factor B |
| 9402 | 1.78e-13 | 1.04e-16 | 0.69 | 8.30 | 5.7090 | 491.35 | GRAP2 | GRB2 related adaptor protein 2 |
| 22806 | 1.61e-12 | 1.08e-15 | 0.56 | 8.02 | 4.5250 | 524.4 | IKZF3 | IKAROS family zinc finger 3 |
| 4514 | 1.61e-12 | 1.17e-15 | 0.66 | 8.01 | 5.2682 | 973257 | COX3 | cytochrome c oxidase subunit III |
| 4579 | 2.73e-12 | 2.18e-15 | 0.67 | 7.93 | 5.3151 | 65989.37 | TRNY | tRNA-Tyr |
| 3575 | 4.61e-12 | 4.02e-15 | 0.81 | 7.85 | 6.3551 | 304.01 | IL7R | interleukin 7 receptor |
| 4511 | 9.34e-12 | 8.82e-15 | 0.67 | 7.76 | 5.1702 | 72358.19 | TRNC | tRNA-Cys |
| 57595 | 1.61e-11 | 1.64e-14 | 0.74 | 7.68 | 5.7026 | 151.43 | PDZD4 | PDZ domain containing 4 |
| 105376333 | 1.80e-11 | 1.96e-14 | 0.60 | 7.65 | 4.6184 | 1825.52 | LOC105376333 | uncharacterized LOC105376333 |
| 3934 | 6.47e-11 | 7.51e-14 | 0.95 | 7.48 | 7.1088 | 364.24 | LCN2 | lipocalin 2 |
| 6374 | 6.84e-10 | 8.44e-13 | 0.83 | 7.15 | 5.9436 | 88.61 | CXCL5 | C-X-C motif chemokine ligand 5 |
| 3932 | 9.55e-10 | 1.25e-12 | 0.72 | 7.10 | 5.1383 | 184.67 | LCK | LCK proto-oncogene, Src family tyrosine kinase |
| 83888 | 9.67e-10 | 1.35e-12 | 0.71 | 7.09 | 5.0282 | 242.84 | FGFBP2 | fibroblast growth factor binding protein 2 |
| 4513 | 9.67e-10 | 1.40e-12 | 0.65 | 7.08 | 4.5795 | 474435.28 | COX2 | cytochrome c oxidase subunit II |
| 7049 | 2.13e-09 | 3.40e-12 | 0.56 | 6.96 | 3.8800 | 348.17 | TGFBR3 | transforming growth factor beta receptor 3 |
| 6352 | 2.13e-09 | 3.34e-12 | 0.72 | 6.96 | 5.0378 | 979.12 | CCL5 | C-C motif chemokine ligand 5 |
| 923 | 3.68e-09 | 6.15e-12 | 0.78 | 6.88 | 5.3526 | 96.99 | CD6 | CD6 molecule |
| 800 | 3.79e-09 | 6.60e-12 | 0.91 | 6.87 | 6.2487 | 138.44 | CALD1 | caldesmon 1 |

| | | | | | | | | |
|---|---|---|---|---|---|---|---|---|
| 7535 | 4.22e-09 | 7.65e-12 | 0.54 | 6.84 | 3.7253 | 633.74 | ZAP70 | zeta chain of T cell receptor associated protein kinase 70 |
| 9235 | 4.44e-09 | 8.38e-12 | 0.78 | 6.83 | 5.3229 | 271.94 | IL32 | interleukin 32 |
| 388228 | 1.55e-08 | 3.04e-11 | 0.75 | 6.64 | 4.9571 | 101.06 | SBK1 | SH3 domain binding kinase 1 |
| 5197 | 1.60e-08 | 3.24e-11 | 0.97 | 6.64 | 6.4052 | 65.81 | PF4V1 | platelet factor 4 variant 1 |
| 340205 | 1.83e-08 | 3.85e-11 | 0.93 | 6.61 | 6.1137 | 94.12 | TREML1 | triggering receptor expressed on myeloid cells like 1 |
| 9806 | 1.85e-08 | 4.04e-11 | 0.68 | 6.60 | 4.4550 | 432.84 | SPOCK2 | SPARC (osteonectin), cwcv and kazal like domains proteoglycan 2 |
| 399844 | 1.85e-08 | 4.17e-11 | 0.60 | 6.60 | 3.9782 | 2082.94 | LINC01002 | long intergenic non-protein coding RNA 1002 |
| 4519 | 1.87e-08 | 4.34e-11 | 0.71 | 6.59 | 4.6491 | 612427.62 | CYTB | cytochrome b |
| 9047 | 2.77e-08 | 6.84e-11 | 0.66 | 6.52 | 4.2718 | 170.51 | SH2D2A | SH2 domain containing 2A |
| 4540 | 2.77e-08 | 6.81e-11 | 0.69 | 6.52 | 4.5075 | 641497.04 | ND5 | NADH dehydrogenase subunit 5 |
| 1667 | 3.17e-08 | 8.30e-11 | 1.06 | 6.50 | 6.8993 | 704.86 | DEFA1 | defensin alpha 1 |
| 728358 | 3.17e-08 | 8.30e-11 | 1.06 | 6.50 | 6.8993 | 704.86 | DEFA1B | defensin alpha 1B |
| 83699 | 3.22e-08 | 8.64e-11 | 0.73 | 6.49 | 4.7208 | 173.77 | SH3BGRL2 | SH3 domain binding glutamate rich protein like 2 |
| 5335 | 3.42e-08 | 9.45e-11 | 0.62 | 6.48 | 4.0306 | 203.92 | PLCG1 | phospholipase C gamma 1 |
| 107984360 | 3.68e-08 | 1.04e-10 | 1.01 | 6.46 | 6.4975 | 67.89 | LOC107984360 | |
| 4538 | 3.73e-08 | 1.08e-10 | 0.69 | 6.45 | 4.4463 | 939275.04 | ND4 | NADH dehydrogenase subunit 4 |
| 3820 | 4.08e-08 | 1.21e-10 | 0.76 | 6.44 | 4.8871 | 124.97 | KLRB1 | killer cell lectin like receptor B1 |
| 10417 | 5.66e-08 | 1.77e-10 | 0.59 | 6.38 | 3.7395 | 430.09 | SPON2 | spondin 2 |
| 915 | 5.66e-08 | 1.75e-10 | 0.93 | 6.38 | 5.9533 | 63.29 | CD3D | CD3 delta subunit of T-cell receptor complex |
| 10666 | 6.27e-08 | 2.00e-10 | 0.74 | 6.36 | 4.7147 | 503.28 | CD226 | CD226 molecule |
| 919 | 1.05e-07 | 3.42e-10 | 0.72 | 6.28 | 4.5127 | 332.77 | CD247 | CD247 molecule |
| 101928512 | 1.11e-07 | 3.71e-10 | 0.68 | 6.27 | 4.2679 | 127.67 | LOC101928512 | uncharacterized LOC101928512 |
| 8784 | 1.16e-07 | 4.05e-10 | 0.86 | 6.25 | 5.3934 | 129.67 | TNFRSF18 | TNF receptor superfamily member 18 |
| 3001 | 1.16e-07 | 4.01e-10 | 0.82 | 6.25 | 5.1487 | 272.2 | GZMA | granzyme A |

| | | | | | | | | |
|---|---|---|---|---|---|---|---|---|
| 30009 | 1.16e-07 | 4.12e-10 | 0.69 | 6.25 | 4.3091 | 289.63 | TBX21 | T-box transcription factor 21 |
| 171558 | 1.48e-07 | 5.36e-10 | 1.04 | 6.21 | 6.4553 | 74.34 | PTCRA | pre T cell antigen receptor alpha |
| 219670 | 1.95e-07 | 7.20e-10 | 1.12 | 6.16 | 6.9115 | 53.56 | ENKUR | enkurin, TRPC channel interacting protein |
| 116987 | 2.02e-07 | 7.64e-10 | 0.78 | 6.15 | 4.8038 | 82.03 | AGAP1 | ArfGAP with GTPase domain, ankyrin repeat and PH domain 1 |
| 84433 | 2.21e-07 | 8.51e-10 | 0.58 | 6.14 | 3.5854 | 494.22 | CARD11 | caspase recruitment domain family member 11 |
| 3702 | 2.43e-07 | 9.52e-10 | 0.80 | 6.12 | 4.9201 | 181.27 | ITK | IL2 inducible T cell kinase |
| 7504 | 2.63e-07 | 1.05e-09 | 1.01 | 6.10 | 6.1531 | 72.04 | XK | X-linked Kx blood group antigen, Kell and VPS13A binding protein |
| 2815 | 3.03e-07 | 1.23e-09 | 0.98 | 6.08 | 5.9768 | 82.38 | GP9 | glycoprotein IX platelet |
| 9848 | 3.25e-07 | 1.34e-09 | 0.79 | 6.06 | 4.8097 | 87.75 | MFAP3L | microfibril associated protein 3 like |
| 84281 | 3.62e-07 | 1.53e-09 | 0.75 | 6.04 | 4.5399 | 256.17 | C2orf88 | chromosome 2 open reading frame 88 |
| 916 | 4.20e-07 | 1.80e-09 | 0.89 | 6.01 | 5.3772 | 118.84 | CD3E | CD3 epsilon subunit of T-cell receptor complex |
| 8631 | 4.37e-07 | 1.90e-09 | 0.67 | 6.01 | 4.0193 | 126.18 | SKAP1 | src kinase associated phosphoprotein 1 |
| 3493 | 4.44e-07 | 1.97e-09 | 0.85 | 6.00 | 5.0916 | 76.01 | IGHA1 | immunoglobulin heavy constant alpha 1 |
| 286 | 6.56e-07 | 2.95e-09 | 0.87 | 5.93 | 5.1352 | 90.7 | ANK1 | ankyrin 1 |
| 55287 | 7.24e-07 | 3.31e-09 | 1.08 | 5.92 | 6.4090 | 57.91 | TMEM40 | transmembrane protein 40 |
| 113219467 | 7.37e-07 | 3.42e-09 | 0.76 | 5.91 | 4.5044 | 76179.7 | MIR12136 | microRNA 12136 |
| 321 | 7.39e-07 | 3.49e-09 | 0.93 | 5.91 | 5.4982 | 147.92 | APBA2 | amyloid beta precursor protein binding family A member 2 |
| 4574 | 7.85e-07 | 3.76e-09 | 0.77 | 5.89 | 4.5471 | 35610.06 | TRNS1 | tRNA-Ser |
| 2781 | 8.06e-07 | 3.92e-09 | 0.83 | 5.89 | 4.8570 | 70.27 | GNAZ | G protein subunit alpha z |
| 53637 | 8.09e-07 | 3.99e-09 | 0.90 | 5.88 | 5.3183 | 170.03 | S1PR5 | sphingosine-1-phosphate receptor 5 |
| 3002 | 8.70e-07 | 4.36e-09 | 0.79 | 5.87 | 4.6264 | 378.29 | GZMB | granzyme B |
| 925 | 8.77e-07 | 4.57e-09 | 1.01 | 5.86 | 5.9427 | 145.77 | CD8A | CD8a molecule |
| 2017 | 8.77e-07 | 4.47e-09 | 0.84 | 5.87 | 4.9530 | 125.5 | CTTN | cortactin |
| 27087 | 8.77e-07 | 4.58e-09 | 0.73 | 5.86 | 4.3039 | 92.01 | B3GAT1 | beta-1,3-glucuronyltransferase 1 |
| 4563 | 9.39e-07 | 4.97e-09 | 0.84 | 5.85 | 4.9028 | 850.27 | TRNG | tRNA-Gly |

| | | | | | | | | |
|---|---|---|---|---|---|---|---|---|
| 2039 | 9.40e-07 | 5.05e-09 | 0.83 | 5.85 | 4.8690 | 158.75 | DMTN | dematin actin binding protein |
| 940 | 1.01e-06 | 5.48e-09 | 1.31 | 5.83 | 7.6261 | 62.42 | CD28 | CD28 molecule |
| 5551 | 1.20e-06 | 6.60e-09 | 0.64 | 5.80 | 3.7325 | 814.54 | PRF1 | perforin 1 |
| 9580 | 1.51e-06 | 8.45e-09 | 0.90 | 5.76 | 5.2089 | 57.92 | SOX13 | SRY-box transcription factor 13 |
| 3824 | 1.63e-06 | 9.33e-09 | 0.76 | 5.74 | 4.3793 | 701.43 | KLRD1 | killer cell lectin like receptor D1 |
| 4157 | 1.63e-06 | 9.32e-09 | 0.72 | 5.74 | 4.1180 | 107.67 | MC1R | melanocortin 1 receptor |
| 23531 | 2.10e-06 | 1.22e-08 | 0.75 | 5.70 | 4.2533 | 356.73 | MMD | monocyte to macrophage differentiation associated |
| 10578 | 2.17e-06 | 1.28e-08 | 0.77 | 5.69 | 4.3968 | 2002.81 | GNLY | granulysin |
| 3690 | 2.18e-06 | 1.30e-08 | 0.77 | 5.69 | 4.3970 | 947.38 | ITGB3 | integrin subunit beta 3 |
| 4646 | 2.19e-06 | 1.32e-08 | 0.78 | 5.68 | 4.4177 | 83.08 | MYO6 | myosin VI |
| 4576 | 2.74e-06 | 1.67e-08 | 0.80 | 5.64 | 4.5201 | 2420 | TRNT | tRNA-Thr |
| 9651 | 3.05e-06 | 1.88e-08 | 0.68 | 5.62 | 3.8051 | 181.4 | PLCH2 | phospholipase C eta 2 |
| 29094 | 3.51e-06 | 2.19e-08 | 0.84 | 5.60 | 4.6699 | 138.21 | LGALSL | galectin like |
| 113218488 | 3.99e-06 | 2.52e-08 | 0.79 | 5.57 | 4.4242 | 1190.1 | MIR10396B | microRNA 10396b |
| 917 | 4.20e-06 | 2.68e-08 | 1.26 | 5.56 | 7.0159 | 43.66 | CD3G | CD3 gamma subunit of T-cell receptor complex |
| 1731 | 4.28e-06 | 2.77e-08 | 0.62 | 5.56 | 3.4521 | 165.85 | SEPTIN1 | septin 1 |
| 100133331 | 4.54e-06 | 2.97e-08 | 0.59 | 5.54 | 3.2556 | 1856.97 | LOC100133331 | replaced by ID 100288069 |
| 107987001 | 5.10e-06 | 3.37e-08 | 0.75 | 5.52 | 4.1566 | 123.26 | FAM27E4 | family with sequence similarity 27 member E4 |
| 105377384 | 5.24e-06 | 3.51e-08 | 1.78 | 5.51 | 9.8185 | 36.96 | LOC105377384 | uncharacterized LOC105377384 |
| 22885 | 5.24e-06 | 3.54e-08 | 1.02 | 5.51 | 5.6055 | 48.88 | ABLIM3 | actin binding LIM protein family member 3 |
| 2812 | 5.95e-06 | 4.06e-08 | 0.99 | 5.49 | 5.4339 | 165.72 | GP1BB | glycoprotein Ib platelet subunit beta |
| 4818 | 7.21e-06 | 4.97e-08 | 0.70 | 5.45 | 3.8226 | 662.86 | NKG7 | natural killer cell granule protein 7 |
| 22981 | 7.48e-06 | 5.21e-08 | 0.98 | 5.44 | 5.3075 | 64.59 | NINL | ninein like |
| 28526 | 8.20e-06 | 5.78e-08 | 0.83 | 5.43 | 4.4807 | 165.56 | TRDC | T cell receptor delta constant |

| | | | | | | | | |
|---|---|---|---|---|---|---|---|---|
| 116173 | 8.31e-06 | 5.91e-08 | 1.19 | 5.42 | 6.4650 | 50.94 | CMTM5 | CKLF like MARVEL transmembrane domain containing 5 |
| 4555 | 9.49e-06 | 6.82e-08 | 0.79 | 5.40 | 4.2726 | 353.74 | TRND | tRNA-Asp |
| 51176 | 9.82e-06 | 7.13e-08 | 1.02 | 5.39 | 5.4696 | 100.03 | LEF1 | lymphoid enhancer binding factor 1 |
| 3560 | 9.90e-06 | 7.26e-08 | 0.81 | 5.38 | 4.3773 | 842.89 | IL2RB | interleukin 2 receptor subunit beta |
| 100506736 | 1.00e-05 | 7.41e-08 | 0.64 | 5.38 | 3.4365 | 213.79 | SLFN12L | schlafen family member 12 like |
| 139105 | 1.24e-05 | 9.26e-08 | 1.42 | 5.34 | 7.5958 | 41.53 | BEND2 | BEN domain containing 2 |
| 100507747 | 1.41e-05 | 1.06e-07 | 0.72 | 5.32 | 3.8251 | 387.43 | C13orf46 | chromosome 13 open reading frame 46 |
| 57121 | 1.57e-05 | 1.19e-07 | 0.90 | 5.29 | 4.7810 | 57.28 | LPAR5 | lysophosphatidic acid receptor 5 |
| 10225 | 1.69e-05 | 1.30e-07 | 0.73 | 5.28 | 3.8536 | 249.02 | CD96 | CD96 molecule |
| 4318 | 1.83e-05 | 1.44e-07 | 1.00 | 5.26 | 5.2383 | 60.45 | MMP9 | matrix metallopeptidase 9 |
| 128611 | 1.83e-05 | 1.44e-07 | 0.70 | 5.26 | 3.6576 | 276.18 | ZNF831 | zinc finger protein 831 |
| 152789 | 1.94e-05 | 1.53e-07 | 1.01 | 5.25 | 5.3183 | 46.41 | JAKMIP1 | janus kinase and microtubule interacting protein 1 |
| 389813 | 2.04e-05 | 1.63e-07 | 0.79 | 5.24 | 4.1191 | 80.25 | AJM1 | apical junction component 1 homolog |
| 100132062 | 2.17e-05 | 1.75e-07 | 0.59 | 5.22 | 3.0796 | 1831.83 | LOC100132062 | uncharacterized LOC100132062 |
| 4068 | 2.35e-05 | 1.91e-07 | 0.98 | 5.21 | 5.0978 | 53.69 | SH2D1A | SH2 domain containing 1A |
| 10316 | 2.41e-05 | 1.99e-07 | 0.71 | 5.20 | 3.7122 | 101.1 | NMUR1 | neuromedin U receptor 1 |
| 2791 | 2.41e-05 | 2.01e-07 | 0.86 | 5.20 | 4.4424 | 208.83 | GNG11 | G protein subunit gamma 11 |
| 342618 | 2.41e-05 | 2.01e-07 | 0.91 | 5.20 | 4.7030 | 103.18 | SLFN14 | schlafen family member 14 |
| 5159 | 2.99e-05 | 2.52e-07 | 0.76 | 5.16 | 3.9102 | 102.47 | PDGFRB | platelet derived growth factor receptor beta |
| 23762 | 3.02e-05 | 2.57e-07 | 0.76 | 5.15 | 3.8987 | 79.18 | OSBP2 | oxysterol binding protein 2 |
| 107987206 | 3.11e-05 | 2.68e-07 | 0.83 | 5.14 | 4.2491 | 3084253.92 | LOC107987206 | replaced by ID 6029 |
| 6029 | 3.11e-05 | 2.68e-07 | 0.83 | 5.14 | 4.2491 | 3084253.42 | RN7SL1 | RNA component of signal recognition particle 7SL1 |
| 4541 | 3.16e-05 | 2.75e-07 | 0.84 | 5.14 | 4.3254 | 185894.86 | ND6 | NADH dehydrogenase subunit 6 |
| 2999 | 3.16e-05 | 2.78e-07 | 0.85 | 5.14 | 4.3693 | 128.86 | GZMH | granzyme H |
| 1901 | 4.07e-05 | 3.61e-07 | 0.92 | 5.09 | 4.6579 | 78.89 | S1PR1 | sphingosine-1-phosphate receptor 1 |

| | | | | | | | | |
|---|---|---|---|---|---|---|---|---|
| 401124 | 4.53e-05 | 4.04e-07 | 0.89 | 5.07 | 4.4885 | 82.71 | DTHD1 | death domain containing 1 |
| 101927854 | 5.07e-05 | 4.56e-07 | 1.89 | 5.04 | 9.5540 | 31.05 | THRB-AS2 | THRB antisense RNA 2 |
| 105378179 | 5.99e-05 | 5.48e-07 | 0.84 | 5.01 | 4.2007 | 4041673.74 | LOC105378179 | uncharacterized LOC105378179 |
| 378706 | 5.99e-05 | 5.48e-07 | 0.84 | 5.01 | 4.2008 | 4041641.81 | RN7SL2 | RNA component of signal recognition particle 7SL2 |
| 387882 | 8.09e-05 | 7.46e-07 | 1.01 | 4.95 | 5.0084 | 135.94 | C12orf75 | chromosome 12 open reading frame 75 |
| 1236 | 8.44e-05 | 7.85e-07 | 1.26 | 4.94 | 6.2013 | 33.87 | CCR7 | C-C motif chemokine receptor 7 |
| 4508 | 8.78e-05 | 8.22e-07 | 0.85 | 4.93 | 4.2116 | 605386.23 | ATP6 | ATP synthase F0 subunit 6 |
| 6775 | 9.32e-05 | 8.80e-07 | 0.67 | 4.92 | 3.2960 | 190.45 | STAT4 | signal transducer and activator of transcription 4 |
| 4753 | 9.57e-05 | 9.10e-07 | 1.74 | 4.91 | 8.5485 | 40.75 | NELL2 | neural EGFL like 2 |
| 2625 | 1.03e-04 | 9.87e-07 | 0.77 | 4.89 | 3.7669 | 137.39 | GATA3 | GATA binding protein 3 |
| 9214 | 1.24e-04 | 1.20e-06 | 0.82 | 4.86 | 3.9582 | 90.93 | FCMR | Fc mu receptor |
| 84886 | 1.27e-04 | 1.23e-06 | 0.75 | 4.85 | 3.6446 | 119.31 | C1orf198 | chromosome 1 open reading frame 198 |
| 347404 | 1.39e-04 | 1.37e-06 | 0.99 | 4.83 | 4.8007 | 48.83 | LANCL3 | LanC like family member 3 |
| 8654 | 1.45e-04 | 1.43e-06 | 0.70 | 4.82 | 3.3488 | 165.13 | PDE5A | phosphodiesterase 5A |
| 4568 | 1.49e-04 | 1.48e-06 | 0.94 | 4.81 | 4.5051 | 868.52 | TRNL2 | tRNA-Leu |
| 2113 | 1.49e-04 | 1.50e-06 | 0.70 | 4.81 | 3.3880 | 955.33 | ETS1 | ETS proto-oncogene 1, transcription factor |
| 109864279 | 1.52e-04 | 1.53e-06 | 0.83 | 4.81 | 3.9706 | 538366.19 | RNA45SN2 | RNA, 45S pre-ribosomal N2 |
| 114804 | 1.62e-04 | 1.64e-06 | 1.20 | 4.79 | 5.7492 | 36.05 | RNF157 | ring finger protein 157 |
| 100131997 | 1.67e-04 | 1.71e-06 | 0.90 | 4.78 | 4.2977 | 114.65 | FAM27E3 | family with sequence similarity 27 member E3 |
| 2811 | 1.67e-04 | 1.72e-06 | 0.74 | 4.78 | 3.5548 | 200.93 | GP1BA | glycoprotein Ib platelet subunit alpha |
| 100008589 | 2.15e-04 | 2.23e-06 | 0.58 | 4.73 | 2.7227 | 1773.16 | RNA28SN5 | RNA, 28S ribosomal N5 |
| 5727 | 2.75e-04 | 2.88e-06 | 0.87 | 4.68 | 4.0578 | 63.79 | PTCH1 | patched 1 |
| 914 | 3.16e-04 | 3.33e-06 | 0.98 | 4.65 | 4.5406 | 154.04 | CD2 | CD2 molecule |

| | | | | | | | | |
|---|---|---|---|---|---|---|---|---|
| 4509 | 3.46e-04 | 3.67e-06 | 0.81 | 4.63 | 3.7444 | 70464.84 | ATP8 | ATP synthase F0 subunit 8 |
| 100233209 | 3.64e-04 | 3.88e-06 | 0.94 | 4.62 | 4.3254 | 57.35 | PCED1B-AS1 | PCED1B antisense RNA 1 |
| 8555 | 3.67e-04 | 3.95e-06 | 1.16 | 4.61 | 5.3280 | 32.12 | CDC14B | cell division cycle 14B |
| 109864272 | 3.78e-04 | 4.09e-06 | 0.58 | 4.61 | 2.6636 | 1416.82 | RNA28SN4 | RNA, 28S ribosomal N4 |
| 109864282 | 4.00e-04 | 4.36e-06 | 0.58 | 4.59 | 2.6780 | 1379.87 | RNA28SN2 | RNA, 28S ribosomal N2 |
| 1292 | 4.05e-04 | 4.43e-06 | 1.01 | 4.59 | 4.6293 | 484.55 | COL6A2 | collagen type VI alpha 2 chain |
| 80739 | 4.31e-04 | 4.75e-06 | 0.91 | 4.58 | 4.1749 | 571.27 | MPIG6B | megakaryocyte and platelet inhibitory receptor G6b |
| 105379857 | 4.54e-04 | 5.04e-06 | 0.84 | 4.56 | 3.8449 | 58.53 | LOC105379857 | replaced by ID 642819 |
| 4638 | 4.54e-04 | 5.08e-06 | 0.91 | 4.56 | 4.1321 | 130.67 | MYLK | myosin light chain kinase |
| 6678 | 4.66e-04 | 5.24e-06 | 0.82 | 4.55 | 3.7413 | 360.13 | SPARC | secreted protein acidic and cysteine rich |
| 8436 | 5.11e-04 | 5.82e-06 | 0.90 | 4.53 | 4.0978 | 545.74 | CAVIN2 | caveolae associated protein 2 |
| 378707 | 5.11e-04 | 5.86e-06 | 0.76 | 4.53 | 3.4260 | 338691.09 | RN7SL3 | RNA component of signal recognition particle 7SL3 |
| 105372446 | 5.11e-04 | 5.86e-06 | 1.06 | 4.53 | 4.7926 | 39.8 | LIM2-AS1 | LIM2 and SIGLEC10 antisense RNA 1 |
| 106632264 | 5.16e-04 | 5.95e-06 | 0.59 | 4.53 | 2.6802 | 1688.58 | RNA28SN1 | RNA, 28S ribosomal N1 |
| 5730 | 5.73e-04 | 6.66e-06 | 0.85 | 4.50 | 3.8484 | 74.51 | PTGDS | prostaglandin D2 synthase |
| 348378 | 6.02e-04 | 7.04e-06 | 1.00 | 4.49 | 4.5061 | 49.36 | SHISAL2A | shisa like 2A |
| 81606 | 6.29e-04 | 7.40e-06 | 0.74 | 4.48 | 3.3265 | 112.26 | LBH | LBH regulator of WNT signaling pathway |
| 79616 | 6.35e-04 | 7.52e-06 | 1.03 | 4.48 | 4.6155 | 38.45 | CCNJL | cyclin J like |
| 4535 | 6.37e-04 | 7.58e-06 | 0.86 | 4.48 | 3.8383 | 713934.18 | ND1 | NADH dehydrogenase subunit 1 |
| 5577 | 6.62e-04 | 7.93e-06 | 0.77 | 4.47 | 3.4346 | 474.72 | PRKAR2B | protein kinase cAMP-dependent type II regulatory subunit beta |
| 4564 | 6.70e-04 | 8.08e-06 | 0.93 | 4.46 | 4.1357 | 1883.31 | TRNH | tRNA-His |
| 107986461 | 6.71e-04 | 8.13e-06 | 1.32 | 4.46 | 5.9042 | 26.15 | LOC107986461 | uncharacterized LOC107986461 |

| | | | | | | | | |
|---|---|---|---|---|---|---|---|---|
| 4566 | 7.44e-04 | 9.07e-06 | 0.89 | 4.44 | 3.9404 | 1543.83 | TRNK | tRNA-Lys |
| 9886 | 7.46e-04 | 9.16e-06 | 0.78 | 4.44 | 3.4675 | 92.92 | RHOBTB1 | Rho related BTB domain containing 1 |
| 728262 | 8.18e-04 | 1.01e-05 | 0.59 | 4.41 | 2.6184 | 2010.11 | FAM157A | family with sequence similarity 157 member A |
| 4550 | 8.18e-04 | 1.02e-05 | 0.90 | 4.41 | 3.9566 | 262.81 | RNR2 | l-rRNA |
| 8530 | 8.88e-04 | 1.11e-05 | 0.71 | 4.39 | 3.1021 | 207.99 | CST7 | cystatin F |
| 6095 | 9.03e-04 | 1.13e-05 | 0.75 | 4.39 | 3.3120 | 244.43 | RORA | RAR related orphan receptor A |
| 85379 | 9.15e-04 | 1.16e-05 | 1.22 | 4.39 | 5.3521 | 27.68 | KIAA1671 | KIAA1671 |
| 54843 | 9.23e-04 | 1.17e-05 | 0.71 | 4.38 | 3.1244 | 141.13 | SYTL2 | synaptotagmin like 2 |
| 28951 | 9.79e-04 | 1.25e-05 | 0.70 | 4.37 | 3.0415 | 101.4 | TRIB2 | tribbles pseudokinase 2 |
| 3003 | 1.03e-03 | 1.33e-05 | 1.34 | 4.36 | 5.8517 | 76.63 | GZMK | granzyme K |
| 100500862 | 1.14e-03 | 1.47e-05 | 0.93 | 4.33 | 4.0309 | 11787.95 | MIR3648-1 | microRNA 3648-1 |
| 103504731 | 1.15e-03 | 1.49e-05 | 0.93 | 4.33 | 4.0231 | 23318.02 | MIR3648-2 | microRNA 3648-2 |
| 100289124 | 1.25e-03 | 1.64e-05 | 1.10 | 4.31 | 4.7453 | 54.17 | FAM27E2 | family with sequence similarity 27 member E2 |
| 50852 | 1.28e-03 | 1.68e-05 | 1.82 | 4.30 | 7.8346 | 25.64 | TRAT1 | T cell receptor associated transmembrane adaptor 1 |
| 149628 | 1.30e-03 | 1.71e-05 | 0.74 | 4.30 | 3.1719 | 148.48 | PYHIN1 | pyrin and HIN domain family member 1 |
| 154075 | 1.30e-03 | 1.76e-05 | 0.84 | 4.29 | 3.6147 | 206.39 | SAMD3 | sterile alpha motif domain containing 3 |
| 23224 | 1.30e-03 | 1.75e-05 | 0.68 | 4.29 | 2.8986 | 1317.56 | SYNE2 | spectrin repeat containing nuclear envelope protein 2 |
| 9437 | 1.30e-03 | 1.74e-05 | 0.91 | 4.30 | 3.9240 | 76.34 | NCR1 | natural cytotoxicity triggering receptor 1 |
| 113218501 | 1.30e-03 | 1.76e-05 | 1.04 | 4.29 | 4.4554 | 268.18 | MIR10396A | microRNA 10396a |
| 4536 | 1.36e-03 | 1.85e-05 | 0.89 | 4.28 | 3.8011 | 734832.3 | ND2 | NADH dehydrogenase subunit 2 |
| 343413 | 1.37e-03 | 1.87e-05 | 0.81 | 4.28 | 3.4488 | 73.61 | FCRL6 | Fc receptor like 6 |
| 6403 | 1.46e-03 | 2.00e-05 | 0.88 | 4.27 | 3.7668 | 135.52 | SELP | selectin P |
| 9254 | 1.69e-03 | 2.34e-05 | 0.88 | 4.23 | 3.7231 | 57.73 | CACNA2D2 | calcium voltage-gated channel auxiliary subunit alpha2delta 2 |

| | | | | | | | | |
|---|---|---|---|---|---|---|---|---|
| 8277 | 1.69e-03 | 2.34e-05 | 1.21 | 4.23 | 5.1177 | 51.12 | TKTL1 | transketolase like 1 |
| 6043 | 1.81e-03 | 2.53e-05 | 0.94 | -4.21 | -3.9531 | 122.45 | SNORA63 | small nucleolar RNA, H/ACA box 63 |
| 9953 | 1.82e-03 | 2.55e-05 | 0.84 | 4.21 | 3.5489 | 94.43 | HS3ST3B1 | heparan sulfate-glucosamine 3-sulfotransferase 3B1 |
| 2920 | 1.85e-03 | 2.60e-05 | 0.64 | -4.21 | -2.6869 | 371.47 | CXCL2 | C-X-C motif chemokine ligand 2 |
| 28639 | 1.86e-03 | 2.63e-05 | 0.95 | 4.20 | 4.0018 | 55.35 | TRBC1 | T cell receptor beta constant 1 |
| 8807 | 1.88e-03 | 2.67e-05 | 0.69 | 4.20 | 2.9035 | 136.42 | IL18RAP | interleukin 18 receptor accessory protein |
| 5583 | 1.88e-03 | 2.69e-05 | 0.60 | 4.20 | 2.5272 | 810.23 | PRKCH | protein kinase C eta |
| 100128731 | 1.91e-03 | 2.75e-05 | 0.59 | 4.19 | 2.4621 | 589.79 | OST4 | oligosaccharyltransferase complex subunit 4, non-catalytic |
| 100130231 | 1.95e-03 | 2.81e-05 | 1.03 | 4.19 | 4.3111 | 108.49 | LINC00861 | long intergenic non-protein coding RNA 861 |
| 666 | 2.02e-03 | 2.93e-05 | 0.94 | 4.18 | 3.9099 | 43.61 | BOK | BCL2 family apoptosis regulator BOK |
| 143872 | 2.09e-03 | 3.05e-05 | 1.18 | 4.17 | 4.9206 | 31.45 | ARHGAP42 | Rho GTPase activating protein 42 |
| 1793 | 2.14e-03 | 3.14e-05 | 0.87 | -4.16 | -3.5991 | 192.46 | DOCK1 | dedicator of cytokinesis 1 |
| 84131 | 2.33e-03 | 3.43e-05 | 0.61 | 4.14 | 2.5401 | 289.46 | CEP78 | centrosomal protein 78 |
| 259215 | 2.48e-03 | 3.68e-05 | 1.24 | 4.13 | 5.1340 | 28.73 | LY6G6F | lymphocyte antigen 6 family member G6F |
| 51348 | 2.48e-03 | 3.69e-05 | 0.95 | 4.13 | 3.8998 | 150 | KLRF1 | killer cell lectin like receptor F1 |
| 81563 | 2.59e-03 | 3.87e-05 | 0.72 | 4.12 | 2.9448 | 137.04 | C1orf21 | chromosome 1 open reading frame 21 |
| 105377806 | 2.68e-03 | 4.03e-05 | 0.85 | 4.11 | 3.4713 | 109.15 | LOC105377806 | |
| 4050 | 2.77e-03 | 4.18e-05 | 0.66 | 4.10 | 2.6877 | 138.21 | LTB | lymphotoxin beta |
| 81794 | 2.98e-03 | 4.53e-05 | 0.65 | 4.08 | 2.6647 | 253.96 | ADAMTS10 | ADAM metallopeptidase with thrombospondin type 1 motif 10 |
| 339541 | 3.11e-03 | 4.75e-05 | 1.00 | -4.07 | -4.0516 | 148.54 | ARMH1 | armadillo like helical domain containing 1 |
| 58486 | 3.25e-03 | 4.97e-05 | 0.72 | -4.06 | -2.9294 | 120.56 | ZBED5 | zinc finger BED-type containing 5 |
| 3693 | 3.59e-03 | 5.52e-05 | 0.83 | 4.03 | 3.3455 | 79.38 | ITGB5 | integrin subunit beta 5 |
| 8787 | 3.68e-03 | 5.69e-05 | 0.93 | 4.03 | 3.7281 | 42.63 | RGS9 | regulator of G protein signaling 9 |

| | | | | | | | | |
|---|---|---|---|---|---|---|---|---|
| 10158 | 3.90e-03 | 6.09e-05 | 1.88 | 4.01 | 7.5241 | 20.9 | PDZK1IP1 | PDZK1 interacting protein 1 |
| 57732 | 3.90e-03 | 6.07e-05 | 0.68 | 4.01 | 2.7084 | 117.73 | ZFYVE28 | zinc finger FYVE-type containing 28 |
| 22914 | 3.90e-03 | 6.12e-05 | 0.94 | 4.01 | 3.7704 | 103.95 | KLRK1 | killer cell lectin like receptor K1 |
| 54796 | 3.98e-03 | 6.27e-05 | 1.01 | 4.00 | 4.0214 | 46.94 | BNC2 | basonuclin 2 |
| 101929531 | 4.34e-03 | 6.87e-05 | 1.33 | 3.98 | 5.2740 | 22.25 | LINC01871 | long intergenic non-protein coding RNA 1871 |
| 79993 | 4.55e-03 | 7.23e-05 | 0.89 | 3.97 | 3.5273 | 135.06 | ELOVL7 | ELOVL fatty acid elongase 7 |
| 93010 | 4.56e-03 | 7.29e-05 | 0.98 | 3.97 | 3.9048 | 37.06 | B3GNT7 | UDP-GlcNAc:betaGal beta-1,3-N-acetylglucosaminyltransferase 7 |
| 864 | 4.57e-03 | 7.32e-05 | 0.60 | 3.97 | 2.3704 | 1024.65 | RUNX3 | RUNX family transcription factor 3 |
| 4900 | 4.76e-03 | 7.67e-05 | 0.68 | 3.95 | 2.6853 | 1430.52 | NRGN | neurogranin |
| 109910382 | 5.10e-03 | 8.25e-05 | 0.55 | 3.94 | 2.1542 | 1042.18 | RNA28SN3 | RNA, 28S ribosomal N3 |
| 105379807 | 5.44e-03 | 8.87e-05 | 1.17 | 3.92 | 4.5978 | 38.71 | LOC105379807 | uncharacterized LOC105379807 |
| 92591 | 5.44e-03 | 8.88e-05 | 0.87 | 3.92 | 3.4114 | 171.51 | ASB16 | ankyrin repeat and SOCS box containing 16 |
| 124602 | 5.65e-03 | 9.27e-05 | 1.11 | 3.91 | 4.3402 | 44.07 | KIF19 | kinesin family member 19 |
| 9289 | 5.80e-03 | 9.56e-05 | 0.68 | 3.90 | 2.6365 | 465.08 | ADGRG1 | adhesion G protein-coupled receptor G1 |
| 117157 | 5.92e-03 | 9.81e-05 | 0.99 | 3.90 | 3.8584 | 117.34 | SH2D1B | SH2 domain containing 1B |
| 54758 | 5.98e-03 | 9.95e-05 | 0.57 | 3.89 | 2.2291 | 299.45 | KLHDC4 | kelch domain containing 4 |
| 760 | 6.08e-03 | 1.01e-04 | 0.82 | 3.89 | 3.1916 | 123.83 | CA2 | carbonic anhydrase 2 |
| 3823 | 6.16e-03 | 1.03e-04 | 0.92 | 3.88 | 3.5710 | 59.19 | KLRC3 | killer cell lectin like receptor C3 |
| 5874 | 6.32e-03 | 1.06e-04 | 0.87 | 3.88 | 3.3555 | 240.25 | RAB27B | RAB27B, member RAS oncogene family |
| 60509 | 6.62e-03 | 1.12e-04 | 0.61 | 3.86 | 2.3535 | 382.95 | AGBL5 | AGBL carboxypeptidase 5 |
| 4573 | 6.62e-03 | 1.12e-04 | 0.95 | 3.86 | 3.6809 | 306.24 | TRNR | tRNA-Arg |
| 2274 | 6.69e-03 | 1.14e-04 | 0.88 | 3.86 | 3.4101 | 48.27 | FHL2 | four and a half LIM domains 2 |
| 11098 | 6.80e-03 | 1.16e-04 | 0.91 | 3.85 | 3.5156 | 51.9 | PRSS23 | serine protease 23 |
| 4569 | 6.97e-03 | 1.20e-04 | 0.89 | 3.85 | 3.4226 | 682.49 | TRNM | tRNA-Met |

| | | | | | | | | |
|---|---|---|---|---|---|---|---|---|
| 4537 | 7.23e-03 | 1.25e-04 | 0.92 | 3.84 | 3.5099 | 157586.17 | ND3 | NADH dehydrogenase subunit 3 |
| 4539 | 7.81e-03 | 1.35e-04 | 0.93 | 3.82 | 3.5386 | 139355.36 | ND4L | NADH dehydrogenase subunit 4L |
| 23043 | 8.03e-03 | 1.40e-04 | 0.56 | 3.81 | 2.1283 | 380.44 | TNIK | TRAF2 and NCK interacting kinase |
| 10125 | 8.03e-03 | 1.40e-04 | 0.71 | 3.81 | 2.6914 | 163.54 | RASGRP1 | RAS guanyl releasing protein 1 |
| 26030 | 8.07e-03 | 1.42e-04 | 0.53 | 3.81 | 2.0257 | 470.15 | PLEKHG3 | pleckstrin homology and RhoGEF domain containing G3 |
| 6932 | 8.10e-03 | 1.43e-04 | 0.82 | 3.80 | 3.1000 | 387.11 | TCF7 | transcription factor 7 |
| 9495 | 8.39e-03 | 1.49e-04 | 0.93 | 3.79 | 3.5344 | 48.48 | AKAP5 | A-kinase anchoring protein 5 |
| 107987026 | 8.62e-03 | 1.53e-04 | 0.93 | 3.79 | 3.5297 | 95.78 | LOC107987026 | uncharacterized LOC107987026 |
| 109864271 | 8.86e-03 | 1.58e-04 | 0.90 | 3.78 | 3.4098 | 426294.63 | RNA45SN4 | RNA, 45S pre-ribosomal N4 |
| 10123 | 9.12e-03 | 1.64e-04 | 0.58 | 3.77 | 2.1911 | 1100.53 | ARL4C | ADP ribosylation factor like GTPase 4C |
| 100528032 | 9.52e-03 | 1.71e-04 | 0.93 | 3.76 | 3.5015 | 120.66 | KLRC4-KLRK1 | KLRC4-KLRK1 readthrough |
| 820 | 1.03e-02 | 1.87e-04 | 1.21 | 3.74 | 4.5029 | 32.12 | CAMP | cathelicidin antimicrobial peptide |
| 83988 | 1.03e-02 | 1.87e-04 | 0.79 | 3.74 | 2.9563 | 63.94 | NCALD | neurocalcin delta |
| 11278 | 1.07e-02 | 1.95e-04 | 0.75 | 3.73 | 2.8096 | 242.64 | KLF12 | KLF transcription factor 12 |
| 57211 | 1.08e-02 | 1.97e-04 | 1.88 | -3.72 | -7.0025 | 68.04 | ADGRG6 | adhesion G protein-coupled receptor G6 |
| 387509 | 1.11e-02 | 2.04e-04 | 1.05 | 3.71 | 3.9001 | 32.45 | GPR153 | G protein-coupled receptor 153 |
| 4575 | 1.11e-02 | 2.04e-04 | 0.98 | 3.71 | 3.6310 | 1251.35 | TRNS2 | tRNA-Ser |
| 55930 | 1.12e-02 | 2.07e-04 | 1.82 | -3.71 | -6.7664 | 65.57 | MYO5C | myosin VC |
| 102724646 | 1.13e-02 | 2.10e-04 | 0.71 | 3.71 | 2.6310 | 165.84 | LOC102724646 | uncharacterized LOC102724646 |
| 129049 | 1.13e-02 | 2.12e-04 | 1.47 | 3.70 | 5.4433 | 22.97 | SGSM1 | small G protein signaling modulator 1 |
| 440823 | 1.25e-02 | 2.34e-04 | 0.72 | 3.68 | 2.6367 | 193.92 | MIAT | myocardial infarction associated transcript |
| 9124 | 1.27e-02 | 2.39e-04 | 0.80 | 3.67 | 2.9344 | 198.16 | PDLIM1 | PDZ and LIM domain 1 |
| 3674 | 1.27e-02 | 2.39e-04 | 0.85 | 3.67 | 3.1086 | 365.67 | ITGA2B | integrin subunit alpha 2b |
| 84879 | 1.31e-02 | 2.49e-04 | 1.64 | -3.66 | -5.9959 | 37.54 | MFSD2A | MFSD2 lysolipid transporter A, lysophospholipid |

| | | | | | | | | |
|---|---|---|---|---|---|---|---|---|
| 23345 | 1.31e-02 | 2.49e-04 | 0.53 | 3.66 | 1.9435 | 2229.53 | SYNE1 | spectrin repeat containing nuclear envelope protein 1 |
| 105376626 | 1.32e-02 | 2.52e-04 | 1.92 | -3.66 | -7.0378 | 27.27 | LOC105376626 | uncharacterized LOC105376626 |
| 54438 | 1.33e-02 | 2.55e-04 | 0.56 | 3.66 | 2.0469 | 310.53 | GFOD1 | glucose-fructose oxidoreductase domain containing 1 |
| 2696 | 1.35e-02 | 2.59e-04 | 1.16 | 3.65 | 4.2418 | 26.77 | GIPR | gastric inhibitory polypeptide receptor |
| 55655 | 1.35e-02 | 2.61e-04 | 1.06 | 3.65 | 3.8808 | 45.33 | NLRP2 | NLR family pyrin domain containing 2 |
| 54855 | 1.41e-02 | 2.74e-04 | 1.11 | 3.64 | 4.0208 | 282.75 | TENT5C | terminal nucleotidyltransferase 5C |
| 161882 | 1.43e-02 | 2.79e-04 | 0.65 | 3.63 | 2.3771 | 129.01 | ZFPM1 | zinc finger protein, FOG family member 1 |
| 26051 | 1.52e-02 | 2.97e-04 | 0.60 | 3.62 | 2.1631 | 192.26 | PPP1R16B | protein phosphatase 1 regulatory subunit 16B |
| 112268284 | 1.52e-02 | 2.97e-04 | 0.73 | 3.62 | 2.6257 | 547.02 | LOC112268284 | |
| 28638 | 1.55e-02 | 3.06e-04 | 0.90 | 3.61 | 3.2520 | 98.06 | TRBC2 | T cell receptor beta constant 2 |
| 3039 | 1.61e-02 | 3.18e-04 | 2.13 | 3.60 | 7.6657 | 3003.76 | HBA1 | hemoglobin subunit alpha 1 |
| 8542 | 1.62e-02 | 3.21e-04 | 0.76 | 3.60 | 2.7336 | 760.21 | APOL1 | apolipoprotein L1 |
| 122416 | 1.75e-02 | 3.49e-04 | 0.76 | 3.58 | 2.7136 | 105.46 | ANKRD9 | ankyrin repeat domain 9 |
| 23336 | 1.75e-02 | 3.50e-04 | 1.06 | 3.57 | 3.7834 | 31.75 | SYNM | synemin |
| 3004 | 1.75e-02 | 3.49e-04 | 1.19 | 3.58 | 4.2615 | 31.8 | GZMM | granzyme M |
| 4571 | 1.80e-02 | 3.62e-04 | 0.89 | 3.57 | 3.1609 | 5876.29 | TRNP | tRNA-Pro |
| 284 | 1.84e-02 | 3.71e-04 | 1.34 | -3.56 | -4.7648 | 124.32 | ANGPT1 | angiopoietin 1 |
| 106633800 | 1.85e-02 | 3.75e-04 | 0.80 | -3.56 | -2.8334 | 164.61 | SNORD133 | small nucleolar RNA, C/D box 133 |
| 59352 | 1.86e-02 | 3.78e-04 | 1.10 | 3.55 | 3.9145 | 51.9 | LGR6 | leucine rich repeat containing G protein-coupled receptor 6 |
| 7293 | 1.89e-02 | 3.85e-04 | 0.99 | 3.55 | 3.5203 | 57.36 | TNFRSF4 | TNF receptor superfamily member 4 |
| 9053 | 1.93e-02 | 3.96e-04 | 1.52 | -3.54 | -5.3839 | 75.24 | MAP7 | microtubule associated protein 7 |
| 3494 | 1.93e-02 | 3.96e-04 | 1.22 | 3.54 | 4.3330 | 30.15 | IGHA2 | immunoglobulin heavy constant alpha 2 (A2m marker) |

| | | | | | | | | |
|---|---|---|---|---|---|---|---|---|
| 3738 | 1.96e-02 | 4.03e-04 | 0.59 | 3.54 | 2.0960 | 401.84 | KCNA3 | potassium voltage-gated channel subfamily A member 3 |
| 494470 | 1.99e-02 | 4.12e-04 | 1.00 | 3.53 | 3.5138 | 62.46 | RNF165 | ring finger protein 165 |
| 139065 | 1.99e-02 | 4.13e-04 | 0.94 | -3.53 | -3.3180 | 76.99 | SLITRK4 | SLIT and NTRK like family member 4 |
| 202020 | 2.09e-02 | 4.35e-04 | 1.92 | -3.52 | -6.7558 | 22.42 | TAPT1-AS1 | TAPT1 antisense RNA 1 (head to head) |
| 5521 | 2.10e-02 | 4.41e-04 | 0.94 | 3.51 | 3.3171 | 51.66 | PPP2R2B | protein phosphatase 2 regulatory subunit Bbeta |
| 256691 | 2.10e-02 | 4.39e-04 | 1.45 | -3.52 | -5.0834 | 113.64 | MAMDC2 | MAM domain containing 2 |
| 23348 | 2.10e-02 | 4.41e-04 | 1.01 | 3.51 | 3.5339 | 53.89 | DOCK9 | dedicator of cytokinesis 9 |
| 103021164 | 2.10e-02 | 4.43e-04 | 2.02 | 3.51 | 7.0951 | 15.17 | CASC21 | cancer susceptibility 21 |
| 84002 | 2.10e-02 | 4.45e-04 | 0.98 | -3.51 | -3.4341 | 150.14 | B3GNT5 | UDP-GlcNAc:betaGal beta-1,3-N-acetylglucosaminyltransferase 5 |
| 6622 | 2.15e-02 | 4.57e-04 | 0.83 | 3.50 | 2.9029 | 160.64 | SNCA | synuclein alpha |
| 2322 | 2.17e-02 | 4.65e-04 | 0.82 | -3.50 | -2.8844 | 205.22 | FLT3 | fms related receptor tyrosine kinase 3 |
| 3040 | 2.17e-02 | 4.62e-04 | 2.22 | 3.50 | 7.7566 | 3930.13 | HBA2 | hemoglobin subunit alpha 2 |
| 6915 | 2.17e-02 | 4.65e-04 | 1.28 | 3.50 | 4.4765 | 34.89 | TBXA2R | thromboxane A2 receptor |
| 100037417 | 2.35e-02 | 5.07e-04 | 0.75 | 3.48 | 2.6199 | 275.03 | DDTL | D-dopachrome tautomerase like |
| 5729 | 2.41e-02 | 5.22e-04 | 0.91 | 3.47 | 3.1667 | 65.68 | PTGDR | prostaglandin D2 receptor |
| 8809 | 2.53e-02 | 5.48e-04 | 0.98 | 3.46 | 3.3904 | 73.66 | IL18R1 | interleukin 18 receptor 1 |
| 10178 | 2.64e-02 | 5.75e-04 | 0.96 | 3.44 | 3.2996 | 49.23 | TENM1 | teneurin transmembrane protein 1 |
| 105369402 | 2.65e-02 | 5.79e-04 | 1.94 | -3.44 | -6.6822 | 21.29 | LOC105369402 | uncharacterized LOC105369402 |
| 652966 | 2.68e-02 | 5.88e-04 | 0.93 | -3.44 | -3.2043 | 118.06 | SNORD10 | small nucleolar RNA, C/D box 10 |
| 10826 | 2.69e-02 | 5.92e-04 | 0.69 | 3.44 | 2.3688 | 210.39 | FAXDC2 | fatty acid hydroxylase domain containing 2 |
| 6080 | 2.84e-02 | 6.27e-04 | 0.74 | -3.42 | -2.5439 | 587.68 | SNORA73A | small nucleolar RNA, H/ACA box 73A |
| 54541 | 2.89e-02 | 6.40e-04 | 0.64 | 3.41 | 2.1916 | 140.83 | DDIT4 | DNA damage inducible transcript 4 |

| | | | | | | | | |
|---|---|---|---|---|---|---|---|---|
| 8320 | 2.95e-02 | 6.55e-04 | 1.12 | 3.41 | 3.8075 | 35.91 | EOMES | eomesodermin |
| 8425 | 2.95e-02 | 6.57e-04 | 0.67 | 3.41 | 2.2964 | 323.27 | LTBP4 | latent transforming growth factor beta binding protein 4 |
| 8510 | 2.96e-02 | 6.64e-04 | 0.81 | 3.40 | 2.7449 | 81.45 | MMP23 B | matrix metallopeptidase 23B |
| 55607 | 2.96e-02 | 6.62e-04 | 1.01 | 3.40 | 3.4504 | 32.61 | PPP1R9 A | protein phosphatase 1 regulatory subunit 9A |
| 4684 | 3.15e-02 | 7.09e-04 | 0.87 | 3.39 | 2.9587 | 221.2 | NCAM1 | neural cell adhesion molecule 1 |
| 158038 | 3.16e-02 | 7.13e-04 | 1.10 | 3.38 | 3.7310 | 27.66 | LINGO2 | leucine rich repeat and Ig domain containing 2 |
| 54797 | 3.20e-02 | 7.24e-04 | 0.80 | 3.38 | 2.7041 | 293.08 | MED18 | mediator complex subunit 18 |
| 54910 | 3.20e-02 | 7.27e-04 | 0.72 | 3.38 | 2.4343 | 114.63 | SEMA4 C | semaphorin 4C |
| 1668 | 3.59e-02 | 8.19e-04 | 2.06 | 3.35 | 6.8952 | 845.14 | DEFA3 | defensin alpha 3 |
| 1005268 33 | 3.65e-02 | 8.36e-04 | 1.09 | 3.34 | 3.6273 | 233.04 | SEPT5-GP1BB | SEPT5-GP1BB readthrough |
| 4570 | 3.65e-02 | 8.37e-04 | 0.81 | 3.34 | 2.6991 | 4503.41 | TRNN | tRNA-Asn |
| 64407 | 3.70e-02 | 8.55e-04 | 0.62 | 3.33 | 2.0712 | 910.06 | RGS18 | regulator of G protein signaling 18 |
| 1116 | 3.70e-02 | 8.57e-04 | 1.17 | 3.33 | 3.9016 | 77.95 | CHI3L1 | chitinase 3 like 1 |
| 4602 | 3.70e-02 | 8.54e-04 | 1.19 | -3.33 | -3.9641 | 309.63 | MYB | MYB proto-oncogene, transcription factor |
| 671 | 3.79e-02 | 8.80e-04 | 0.90 | 3.33 | 3.0079 | 407.29 | BPI | bactericidal permeability increasing protein |
| 1521 | 3.79e-02 | 8.84e-04 | 0.74 | 3.33 | 2.4454 | 352.02 | CTSW | cathepsin W |
| 5243 | 3.95e-02 | 9.24e-04 | 0.88 | 3.31 | 2.9062 | 82.04 | ABCB1 | ATP binding cassette subfamily B member 1 |
| 8690 | 3.99e-02 | 9.36e-04 | 0.90 | -3.31 | -2.9668 | 59.6 | JRKL | JRK like |
| 256380 | 4.00e-02 | 9.40e-04 | 0.90 | 3.31 | 2.9675 | 105.59 | SCML4 | Scm polycomb group protein like 4 |
| 6232 | 4.03e-02 | 9.53e-04 | 0.59 | 3.30 | 1.9478 | 2438.21 | RPS27 | ribosomal protein S27 |
| 26053 | 4.03e-02 | 9.51e-04 | 0.71 | 3.30 | 2.3556 | 184.35 | AUTS2 | activator of transcription and developmental regulator AUTS2 |
| 23365 | 4.07e-02 | 9.67e-04 | 0.72 | 3.30 | 2.3616 | 246.27 | ARHGE F12 | Rho guanine nucleotide exchange factor 12 |
| 57613 | 4.09e-02 | 9.74e-04 | 1.97 | -3.30 | -6.4960 | 18.72 | FAM234 B | family with sequence similarity 234 member B |

| | | | | | | | | |
|---|---|---|---|---|---|---|---|---|
| 652965 | 4.14e-02 | 9.89e-04 | 0.76 | -3.29 | -2.5110 | 441.42 | SNORA48 | small nucleolar RNA, H/ACA box 48 |
| 102724364 | 4.34e-02 | 1.04e-03 | 0.60 | 3.28 | 1.9821 | 212.36 | SEC22B4P | SEC22 homolog B4, pseudogene |
| 4145 | 4.40e-02 | 1.06e-03 | 0.79 | 3.28 | 2.5723 | 216.52 | MATK | megakaryocyte-associated tyrosine kinase |
| 55759 | 4.43e-02 | 1.07e-03 | 0.91 | -3.27 | -2.9844 | 63.19 | WDR12 | WD repeat domain 12 |
| 60312 | 4.52e-02 | 1.09e-03 | 1.08 | 3.27 | 3.5314 | 46.95 | AFAP1 | actin filament associated protein 1 |
| 84628 | 4.63e-02 | 1.12e-03 | 0.69 | 3.26 | 2.2385 | 142.89 | NTNG2 | netrin G2 |
| 4549 | 4.67e-02 | 1.14e-03 | 1.11 | 3.25 | 3.6065 | 148.56 | RNR1 | s-rRNA |
| 100861532 | 4.68e-02 | 1.14e-03 | 0.94 | 3.25 | 3.0486 | 269159.77 | RNA45SN5 | RNA, 45S pre-ribosomal N5 |
| 105379282 | 4.87e-02 | 1.20e-03 | 0.80 | 3.24 | 2.5973 | 156.8 | LOC105379282 | |
| 4092 | 4.87e-02 | 1.19e-03 | 0.73 | 3.24 | 2.3625 | 169.55 | SMAD7 | SMAD family member 7 |
| 79037 | 4.95e-02 | 1.22e-03 | 0.82 | 3.23 | 2.6512 | 75.01 | PVRIG | PVR related immunoglobulin domain containing |
| 144203 | 4.96e-02 | 1.23e-03 | 1.14 | 3.23 | 3.6825 | 26.84 | OVOS2 | alpha-2-macroglobulin like 1 pseudogene |

**Supplemental Table 2.** "Panel Plus" user added genes to the Mouse PanCancer Immune Profiling panel with optimized probe target sequences.

| Customer Identifier | Accession | Position | Target Sequence |
|---|---|---|---|
| abat | NM_001170978.1 | 3561-3660 | GAGCCACAGTGTCATATACAGATACTTCCGCAGGTCCTTAGAGTTCAAAGGGTTTTAATCCAGGACATAAGCAGAAATCGCTCTCTTTAGTGAAGGGAGC |
| Akt1 | NM_001165894.1 | 899-998 | GCCATGAAGATCCTCAAGAAGGAGGTCATCGTCGCCAAGGATGAGGTTGCCCACACGCTTACTGAGAACCGTGTCCTGCAGAACTCTAGGCATCCCTTCC |
| Bad | NM_007522.3 | 1147-1246 | TTCGAGGCCTTAGGAAAAAAAAAGAGGATCGCTGTGTCCCTTTAACAGGGAGAAGAGCTGACGTACAGCTTGAGTCCCTTCCGGTGCGTGCAATAGCCAC |
| bbc3 | NM_133234.1 | 1462-1561 | CCCCAATCCCCATCCATCTCATTGCATAGGTTTAGAGAGAGCACGTGTGACCACTGGCATTCATTTGGGGGGTGGGAGATATTGGCGGAAGCCACCCCAG |
| birc3 | NM_007464.3 | 426-525 | CCCTGTCATCTCACCATGAACATGGTTCAAGACAGCGCCTTTCTAGCCAAGCTGATGAAGAGTGCTGACACCTTTGAGTTGAAGTATGACTTTTCCTGTG |
| blm | NM_001042527.2 | 265-364 | AAAGATGTGAACGTGTCTGAGGCCTTTTCATTCACTGAGTCTCCACTCCACAAACCAAAGCAGCAGGCAAAGATTGAAGGCTTCTTTAAACATTTCCCTG |
| casp9 | NM_015733.4 | 1676-1775 | ATAACTGTCCTGCTAAGATAGGATTTTGAACTGGGGCAGGCTGCTCTTTCCCTTTGGCGATGCAAACATGCTCCTAGCAGCTTTCAGGTTGTAGGGCAAT |
| ctnnb1 | NM_007614.2 | 2976-3075 | TTGGTCGAGGAGTAACAATACAAATGGATTTGGGGAGTGACTCACGCAGTGAAGAATGCACACGAATGGATCACAAGATGGCGTTATCAAACCCTAGCCT |
| dbn1 | NM_019813.4 | 2571-2670 | CGAAATTTAAAACATGGCAATAAATGGCTCGTGGGCTCTGGCTCCCTGGGACCCTTCCCCTTCTCTTTACCCTCGCTGCTTGGTCAGAAGGAATTATCAG |
| ddx52 | NM_030096.2 | 501-600 | AGGAAAAGGTCAACTTCTTTCGGAACAAGCACAAGATACATGTCCAAGGAACTGATCTTCCTGACCCAATTGCTACATTTCAGCAACTTGACCAGGAATA |
| mdm2 | NM_010786.4 | 1665-1764 | GTCATGTTTCACGTGTGCAAAGAAGCTAAAAAAAAGAAACAAGCCCTGCCCAGTGTGCAGACAGCCAATCCAAATGATTGTGCTAACTTACTTCAACTAG |
| pdk4 | NM_013743.2 | 1356-1455 | AGCGGATGACGCCTGACATTTTACGGGATCAAAGTGGGTCTGTGGCATTGCTGCTTCGTGAATGTGTGTGGACTCTAGTTTCCGCAAAACAACGCAACAC |
| psmb5 | NM_011186.1 | 335-434 | TGCAGCTTCTGGGAGCGGTTGTTGGCTCGGCAGTGTCGAATCTATGAGCTTCGCAATAAGGAACGCATCTCGGTCGCAGCAGCCTCCAAACTGCTCGCTA |
| psmc4 | NM_011874.2 | 1147-1246 | TCGACCTGGAAGACTATGTGGCCCGTCCAGATAAGATTTCAGGAGCCGATATCAACTCCATCTGTCAGGAGAGTGGAATGTTGGCTGTCCGTGAGAACCG |
| Ripk1 | NM_009068.3 | 3186-3285 | GCTTTGGCCTTGTTGGCCATTCTGGCACTCATTGGCACTTCATCCTCCTTTGTTGGGCTATCCTGTACTCAGTAGGATATTTGGGAACATTCCTGGCCTC |

| | | | |
|---|---|---|---|
| Ripk3 | NM_019955.1 | 271-370 | CACAGAACATGGAACCATGATGTAGCAGTCAAGATCGTGAACTCGAAGAAGATATCCTGGGAGGTGAAGGCTATGGTTAATCTTCGTAATGAGAACGTTC |
| rock1 | NM_009071.2 | 416-515 | AAGTTGGTTGAACTTGCTTTCCGCTGCGGGCAAGAAGGTATCGTCACAAGTAGCAGCATCATGTCGACTGGGGACAGTTTTGAGACTCGGTTTGAAAAAA |
| slc24a3 | NM_053195.2 | 2271-2370 | CAGGAGAGGGTCCGCTGATGGCAGGAAGGTTTGTTTTGTTTGGGAGTGAGTCCTAGGTTACGGGGCTCAGGGAAATTGTTTAATTTGAGGGGGCGCTTTT |
| tnfsf9 | NM_009404.3 | 293-392 | GCTCTATGGCCTAGTCGCTTTGGTTTTGCTGCTTCTGATCGCCGCCTGTGTTCCTATCTTCACCCGCACCGAGCCTCGGCCAGCGCTCACAATCACCACC |
| ubb | NM_011664.4 | 1324-1423 | TCCTCCGTCTGAGGGGTGGCTATTAATTATTCGGTCTGCATTCCCAGTGGGCAGTGATGGCATTACTCTGCACTCTAGCCACTTGCCCCAATTTAAGTTT |
